# Integrative single-cell analysis of cardiogenesis identifies developmental trajectories and non-coding mutations in congenital heart disease

**DOI:** 10.1101/2022.06.29.498132

**Authors:** Mohamed Ameen, Laksshman Sundaram, Abhimanyu Banerjee, Mengcheng Shen, Soumya Kundu, Surag Nair, Anna Shcherbina, Mingxia Gu, Kitchener D. Wilson, Avyay Varadarajan, Nirmal Vadgama, Akshay Balsubramani, Joseph C. Wu, Jesse Engreitz, Kyle Farh, Ioannis Karakikes, Kevin C Wang, Thomas Quertermous, William Greenleaf, Anshul Kundaje

**Author notes:** These authors contributed equally (Mohamed Ameen, Laksshman Sundaram).

## Abstract

Congenital heart defects, the most common birth disorders, are the clinical manifestation of anomalies in fetal heart development - a complex process involving dynamic spatiotemporal coordination among various precursor cell lineages. This complexity underlies the incomplete understanding of the genetic architecture of congenital heart diseases (CHDs). To define the multi-cellular epigenomic and transcriptional landscape of cardiac cellular development, we generated single-cell chromatin accessibility maps of human fetal heart tissues. We identified eight major differentiation trajectories involving primary cardiac cell types, each associated with dynamic transcription factor (TF) activity signatures. We identified similarities and differences of regulatory landscapes of iPSC-derived cardiac cell types and their *in vivo* counterparts. We interpreted deep learning models that predict cell-type resolved, base-resolution chromatin accessibility profiles from DNA sequence to decipher underlying TF motif lexicons and infer the regulatory impact of non-coding variants. *De novo* mutations predicted to affect chromatin accessibility in arterial endothelium were enriched in CHD cases versus controls. We used CRISPR-based perturbations to validate an enhancer harboring a nominated regulatory CHD mutation, linking it to effects on the expression of a known CHD gene *JARID2*. Together, this work defines the cell-type resolved cis-regulatory sequence determinants of heart development and identifies disruption of cell type-specific regulatory elements as a component of the genetic etiology of CHD.

## Introduction

Congenital heart disease (CHD) is the most common form of developmental birth defect, affecting 1% of live childbirths every year (van der Linde et al. 2011). Approximately one-third of children with CHD have a linked genetic etiology accounting for the disorder. Only 8% of such cases are attributed to mutations in protein-coding gene regions (Zaidi et al. 2013; Homsy et al. 2015; Pediatric Cardiac Genomics Consortium et al. 2013; Jin et al. 2017), strongly suggesting that other causes, including disruption of gene regulation, substantially contribute to the etiology of CHD. However, heart development is a complex symphony of diverse and interacting cell types and phenotypic transformations, making identification of causal non-coding mutations and their impact on gene regulation in disease-relevant cell types challenging (Bruneau 2013).

Organogenesis of the heart begins from two distinct mesodermal cellular progenitors that originate from the primary heart field (PHF) and secondary heart field (SHF). These two mesodermal lineages give rise to three subtypes of heart cells: myocardial cells, epicardial cells, and endocardial cells that later integrate with cells from the neural crest to form a functional human heart (Sylva, van den Hoff, and Moorman 2014; Meilhac and Buckingham 2018; Srivastava 2006). Prior studies that have profiled the single cell transcriptome of the developing human heart have greatly enhanced our understanding of cell types and genes important for cardiogenesis (Suryawanshi et al. 2020; Asp et al. 2019; Miao et al. 2020). However, a comprehensive resource of cell-type resolved cis and trans regulators of gene expression programs across differentiation trajectories in human cardiac development is lacking. This gap in our understanding raises several unresolved questions about transcriptional regulation of cardiogenesis and its dysregulation by non-coding mutations that may cause CHD: 1) What are the dynamic cis-regulatory elements (cREs) and target genes that define cell types and cell state transitions in cardiogenesis? 2) What is the combinatorial lexicon of transcription factor (TF) motifs encoded in these dynamic cREs? 3) Are *de novo* non-coding CHD mutations enriched in cRE landscapes of specific fetal heart cell types? 4) What are the TF binding sites, cREs, and target genes impacted by putative causal non-coding CHD mutations? 5) Which *in vitro* differentiated cellular model systems demonstrably reproduce both the gene expression and chromatin landscape of the *in vivo* developing human heart, thereby enabling functional validation of the regulatory impact of mutations?

Single-cell profiling of chromatin accessibility and gene expression have allowed many of these questions to be addressed in other organ systems, including human hippocampus (Zhong et al. 2020), fetal embryogenesis (Domcke et al. 2020; Cao et al. 2020), human corticogenesis (Domcke et al. 2020; Trevino et al. 2021) and hematopoiesis (Buenrostro et al. 2018; Granja et al. 2019). Cell-type resolved maps of chromatin accessibility provide a window into the dynamic activity of cis- and trans-acting factors, and when combined with gene expression data, can be used to nominate specific TFs, cREs, and regulatory networks associated with cellular state changes. Further, deep learning models that predict chromatin accessibility from DNA sequence have been used to decode TF motif syntax of cREs and nominate putative causal regulatory variants (Avsec, Agarwal, et al. 2021; David R. Kelley, Snoek, and Rinn 2016; D. R. Kelley et al. 2018; D. R. Kelley 2020; J. Zhou and Troyanskaya 2015; Richter et al. 2020; J. Zhou et al. 2018; Trevino et al. 2021). While others have applied such models to tissue-level, bulk cardiac functional genomic data (Richter et al. 2020), the lack of cell-type resolution in these data makes it difficult to decipher cell-type specific effects of variants, especially in rare cell types.

To address these questions, we derived a joint atlas of integrated single-cell data by generating and combining single cell ATAC-seq (scATAC-seq) experiments profiling chromatin landscapes of three primary human fetal heart samples spanning post conception week (PCW) 6, 8 and 19 with published single-cell RNA-seq (scRNA-seq) data at similar developmental timepoints. We deconvolved 20 distinct cell types spanning three progenitor lineages along with neural crest cells and systematically characterized cell-type resolved repertoires of accessible cREs and their putative linked target genes. We trained convolutional neural networks (CNN) that predict cell-type resolved, base-resolution chromatin accessibility profiles from DNA sequence to decipher the dynamic motif lexicon of combinatorial TF binding at all cREs in each cell context (Avsec, Weilert, et al. 2021; Trevino et al. 2021). We adapted an optimal transport algorithm to identify 8 major differentiation trajectories, defining continuous progression of TF activities that promote the formation of primary cell types of the heart (Schiebinger et al. 2019). Using this atlas of cell states representing *in vivo* cardiac development, we compared accessible chromatin landscapes of common cellular model systems comprising major cardiac cell types derived from iPSCs *in vitro*. This comparison revealed substantial epigenomic and transcriptional differences between different model cell-types and their *in vivo* counterparts, except for *in vitro* derived cardiomyocytes and endothelial cells, which showed comparatively high concordance. Finally, we used our deep learning models to prioritize putative causal, non-coding mutations in CHD trios from the Pediatric Cardiac Genomics Consortium (PCGC) (Richter et al. 2020) based on their predicted impact on cell-type specific chromatin accessibility of putative cREs via disruption of TF binding sites. Predicted deleterious mutations in cREs in arterial endothelial cells were enriched (*p-*value = 0.008, odds ratio = 1.7, Fisher’s Exact test) in cases versus healthy controls, thereby revealing one of the predominant cell-types of origin for congenital heart disorders that result from such regulatory mutations. We used CRISPR-based enhancer knockout experiments in *in vitro* differentiated endothelial cells to validate the regulatory impact of a putative cell-type specific enhancer predicted to harbor a deleterious CHD mutation on expression of *JARID2*, an important cell-type specific cardiogenesis gene. Together, these data and models define the cis- and trans-regulatory landscape of the developing human heart across mid-gestation developmental trajectories, elucidate the fidelity of diverse iPSC-to-lineage *in vitro* differentiations, and provide a deep learning framework capable of specifically nominating non-coding *de novo* mutations in candidate cREs predicted to disrupt TF binding, chromatin state and gene expression consistent with causality for CHD.

## Results

### Integrating single-cell ATAC and RNA sequencing data into a unified cell-type resolved regulatory atlas of the developing human heart

To capture chromatin dynamics in different cell populations throughout fetal heart development, we used the Chromium 10X platform to generate scATAC-seq data (Satpathy et al. 2019) from three primary human fetal heart samples at 6-, 8-, and 19-weeks post-conception (PCW) (**Figure 1a)**. We obtained 30,426 high quality scATAC-seq cell barcodes post filtering and quality control (**SFigure 1, Table S1, Methods**). We applied iterative latent semantic indexing (LSI) on accessible chromatin regions to map the cells from all three time points into a multidimensional principal component (PC) space and then to a 2-dimensional uniform manifold approximation and projection (UMAP) representation (Granja et al. 2021; Becht et al. 2018; McInnes et al. 2018). We used the Leiden clustering algorithm to discover and optimize clusters of cells that potentially correspond to distinct cell-types (Traag, Waltman, and van Eck 2019) **(Figure 1b, 1c, SFigure 2, Table S2, Methods)**.

**Figure 1.**
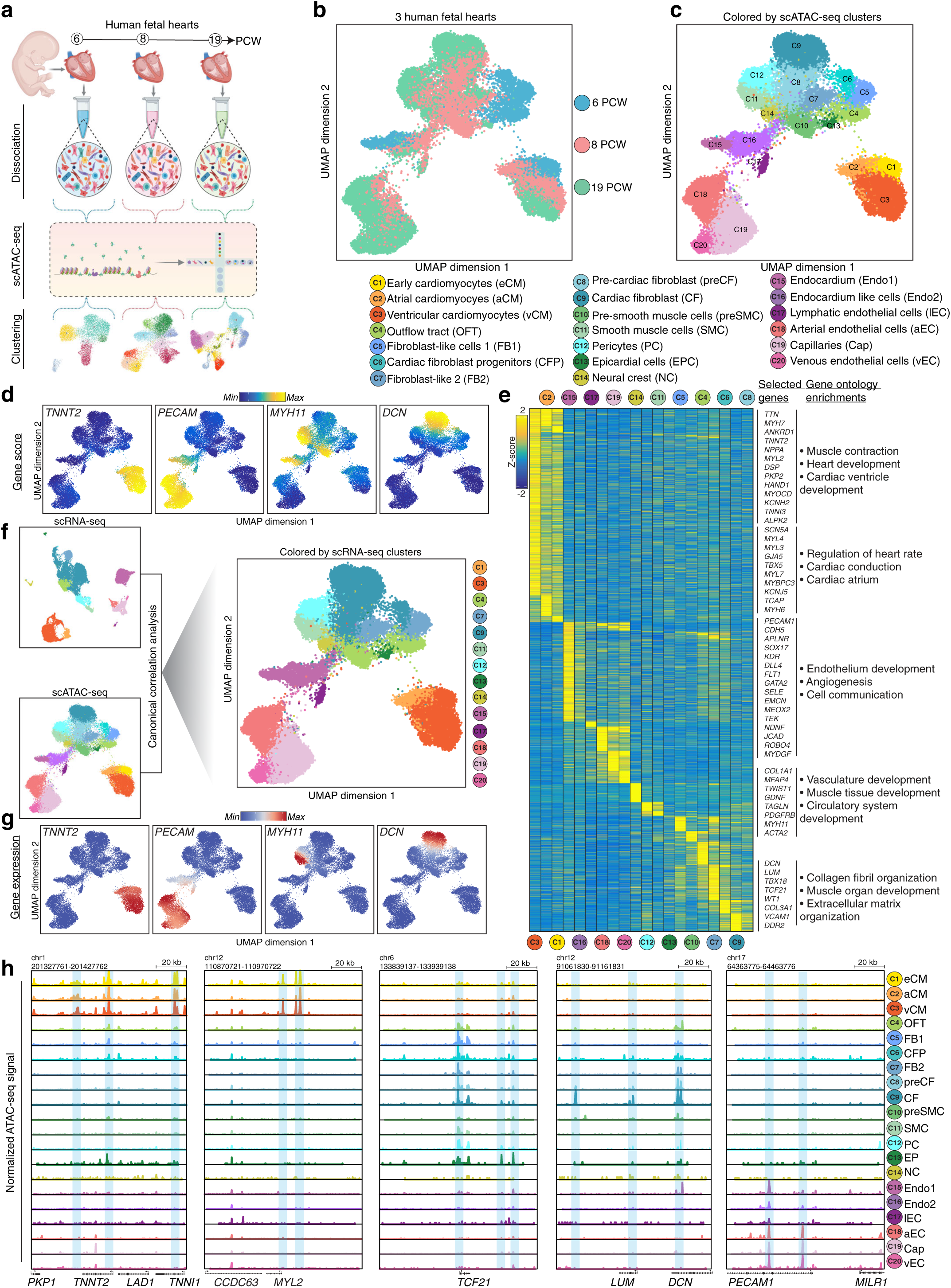
A single-cell epigenomic atlas of the developing human heart. **(a)** Schematic of gestational sample time (post-conception week, PCW) and genome-wide profiling methods represented in this study. **(b)** Uniform Manifold Approximation and Projection (UMAP) of cells based on accessible chromatin regions (scATAC-seq). Cells are colored according to sample gestational time. **(c)** UMAP of cells based on accessible chromatin regions (scATAC-seq). Cells are colored according to cell types identified. **(d)** Single-cell gene accessibility scores (based on scATAC-seq) of *TNNT2, PECAM1, MYH11*, and *DCN*. **(e)** Heatmap of *z*-scores of log_2_(scATAC-seq read counts) in 215,163 *cis*-regulatory elements (cREs) across scATAC-seq cell-type clusters derived from (b). Representative genes with cluster-specific differential gene accessibility scores are shown to the right. Gene ontology enrichments indicate the statistically significant (adjusted *p*-value < 0.005, Gprofiler Fisher’s exact test) cellular processes for genes with differential gene accessibility scores associated with the clusters of cell-type specific cREs. **(f)** UMAPs of scRNA-seq and scATAC-seq cells colored by cluster assignment in their respective data modality, and UMAP of scATAC-seq cells highlighted by complementary scRNA-seq clusters. **(g)** Single-cell gene expression (scRNA-seq) of *TNNT2, PECAM1, MYH11*, and *DCN*. **(h)** Genome tracks of cell-type resolved aggregate scATAC-seq data around the *TNNT2*, *MYL2*, *TCF21*, *DCN/LUM* and *PECAM1* gene loci (left to right). The scale of the tracks (from left to right) range from 0-0.28, 0-0.31, 0-0.18, 0-0.14 and 0-0.2 respectively, in units of fold-enrichment relative to the total number of reads in TSSs per 10k. Highlights indicate the relevant cell type-specific putative enhancers in each gene locus.

We computed chromatin-derived gene accessibility scores by aggregating scATAC-seq reads in each cell weighted by distance from each gene within its cis-regulatory domain (Granja et al. 2021). We deciphered each cluster’s likely cell-type identity based on cluster-specific gene accessibility scores of reference marker genes known to exhibit cell-type specific gene expression (**SFigure 3, Table S3, Methods**). We first defined four different broad precursor lineages - myocardium, epicardium, endocardium, and neural crest (Srivastava 2006) - consistent with the primary heart field (PHF) and secondary heart field (SHF) derived precursor cells in the fetal heart (**Figure 1c**), and then further annotated sub-clusters within these lineages.

Within the myocardial lineage, we found that *TNNT2*, *ACTN2*, and *NKX2-5* had high gene accessibility scores across the early cardiomyocytes (eCM), ventricular cardiomyocytes (vCM), and atrial cardiomyocyte (aCM) clusters (Miao et al. 2020; Cui et al. 2019; Asp et al. 2019; Suryawanshi et al. 2020). *TTN* and *HAND1* activity specifically marked the eCM and vCM clusters while *TBX10*, *NPPA*, and *MYL7* had higher activities in the aCM cluster. The eCM cluster mainly comprised early gestational cells (PCW6) compared to aCM and vCM clusters (**Figure 1d, SFigure 3**).

We observed diverse cell types within the epicardial lineage, which showed varying compositions across different gestational time points. We discovered four cell types at PCW6: cardiac fibroblast progenitors (CFP) with high *WT1*, *TBX18*, and *TCF21* gene accessibility scores, another set of similar cells with both *TBX18* and *TCF21* signal but lacking *WT1* which we call fibroblast-like cells (FB1), and the outflow tract (OFT) like cells with high *PRDM6 (Davis et al. 2006)* and *HOXA3* gene accessibility scores (**SFigure 3**). These OFT cells had low *TCF21* signal and were therefore unlikely to act as cardiac fibroblast progenitors (Acharya et al. 2012). We also found an undifferentiated epicardium cell cluster (EPC) with high *TBX18* and *WT1* signals but lacking *TCF21 (Mikawa and Gourdie 1996; Cai et al. 2008)* (**SFigure 3**). These four clusters appeared to act as progenitor populations for cells arising at later gestational timepoints in the epicardium. PCW8 cells labeled as pre-cardiac fibroblasts (preCF) had high gene accessibility scores for *TCF21* but very low signals for *DCN* and *LUM* (**Figure 1d**, **SFigure 3**). A more mature cardiac fibroblast (CF) population mostly in PCW19 and some in PCW8 cells had high gene accessibility scores for major cardiac fibroblast markers including *DCN*, *LUM*, and *TCF21* (**Figure 1d, SFigure 3**) (Muhl et al. 2020). We defined a second cluster of fibroblast-like cells (FB2) that appeared in PCW8 and PCW19 and exhibited *CNN1* and *COL9A2* gene accessibility scores, but lacked signals for other standard cardiac fibroblast markers (**SFigure 3**). We hypothesize that this cell type, along with FB1, may be related to valvular fibroblasts, but further studies are required to establish this potential relationship. Finally, we defined a cluster of pre-smooth muscle cells (preSMC) with *MYH11*, *PDGFRB*, and *TAGLN* activity but lacking *TCF21* activity (Dobnikar et al. 2018), a cluster of smooth muscle cells (SMC) exhibiting stronger activity for *MYH11* and *PDGFRB* with major contributions from PCW19 and minor contributions from PCW8, and a cluster of pericytes (PC) with activity of *PDGFRB* and *ABCC9* (**Figure 1d, SFigure 3**) (Pham et al. 2021). We also defined a cluster of neural crest (NC) cells with high *TFAP2A* activity (**Figure 1d, SFigure 3**) (W.-D. Wang et al. 2011).

The endocardial cell populations exhibited two distinct phenotypes: one with high *CDH11* activity scores (Endo1) and a smaller population that resembled endocardial-like transitioning cell types (Endo2) (Aird 2007). Arterial endothelial cells (aEC) exhibited high *UNC5B* and *GJA5* gene accessibility score activity. Capillary cells (Cap) were marked by high *CA4*, *APLNR*, and *CD36* gene accessibility scores (**SFigure 3**). Venous endothelial cells (vEC) were marked by high *SELE* and *SELP* gene accessibility score activity, amongst other markers (Kalucka et al. 2020; Vodyanik et al. 2010). In addition to these major endothelial cell types, we also found a sub-population of lymphatic endothelial cells (lEC) exhibiting *LYVE1* gene accessibility score activity (**SFigure 3**) (Podgrabinska et al. 2002). Using these annotated clusters, we identified 215,163 putative cREs as scATAC-seq peak regions over all cell types and timepoints (**Figure 1e**). The clusters were enriched for expected gene ontology (GO) terms associated with cardiac development and cell-type specific attributes (Bruneau 2013) (**Figure 1e, Table S4**).

To understand the correspondence between the chromatin and gene expression landscapes of these cell-types, we analyzed previously published scRNA-seq data from developmental time points that closely match those sampled in our scATAC-seq atlas (Miao et al. 2020; Cui et al. 2019; Asp et al. 2019; Suryawanshi et al. 2020) (**Figure 1f, SFigure 4, Table S5-S6**). Because of the different sources and methods of preparation of cells and scRNA libraries, we harmonized the scRNA-seq data across time points after correcting for batch effects using Harmony (Korsunsky et al. 2019), clustered cells using the Leiden community detection method, and mapped clusters to specific cell-types based on cell-type specific expression of known marker genes (**SFigure 4a,b,c**). Cells from our annotated scATAC-seq atlas were then matched with their nearest neighbor cells in the scRNA-seq atlas using canonical correlation analysis (CCA) (Cusanovich et al. 2018) (**Figure 1f & SFigure 4d**). We found high concordance (accuracy = 74.76%) between the cluster assignments for cells from the scATAC-seq and scRNA-seq data, further supporting our cell type annotations based on chromatin accessibility derived gene accessibility scores (**SFigure 4e**). Examining a subset of cell-type specific marker genes, we found *TNNT2* marking the ventricular cardiomyocytes, *PECAM1* identifying endothelial cells, *CDH11* identifying endocardium, *MYH11* identifying SMC, and *DCN* identifying fibroblasts (Wolf et al. 2019; Ng, Wong, and Tsang 2010) (**Figure 1d,g**). We also observed a strong correlation (**Table S7**) between gene expression from the scRNA-seq data and the gene accessibility scores from the scATAC-seq data across matched nearest-neighbor cells from the two complementary atlases (**Figure 1d,g**), further supporting our annotations.

Next, we used our integrated atlas to examine the complex relationship between the expression of well-known lineage-specific marker genes and chromatin dynamics of their putative cREs. For example, *TNNT2,* a well-known cardiomyocyte marker, exhibited the strongest chromatin accessibility at its promoter and putative distal enhancers, specifically in the three cardiomyocyte clusters (**Figure 1h**). The patterns of accessibility matched the specificity and similarity of expression of *TNNT2* in the same clusters (**SFigure 4c**). In contrast, *MYL2,* a specific marker of ventricular cardiomyocytes, exhibited similar distal chromatin accessibility in the three myocardial lineage clusters, while the promoter was not accessible, and the gene was not expressed, in atrial cardiomyocyte clusters (**Figure 1h, SFigure 4c**), indicating that accessibility of these distal elements may not be sufficient to drive its expression. In the epicardial cell lineage, we observed increasing chromatin accessibility around the *DCN* marker gene through the cardiac fibroblast cell lineage specification (**Figure 1h**) concordant with its gene expression dynamics (**Figure 1g**). We observed analogous dynamics for *PECAM1* in the endocardial lineage. We also observed chromatin state changes consistent with promoter priming for genes in specific cell-types that do not express the associated gene. For instance, the promoter of the developmental gene *TCF21* was accessible in cardiac fibroblast and SMC cell lineages but the gene was expressed only in cardiac fibroblasts and not in SMC (Acharya et al. 2012; Nurnberg et al. 2015) (**Figure 1h, SFigure 4c**). Interestingly, *TCF21* expression is known to be activated in SMC in adults in response to vascular stress (Wirka et al. 2019), promoting cell state changes such as proliferation and migration, consistent with a return to an embryonic-like phenotype for the SMC (Nurnberg et al. 2015). Thus, accessibility of the TSS at the *TCF21* gene may represent adaptive promoter priming (Ma et al. 2020) that allows the gene to rapidly respond to disease-related stress or cellular activation.

### Deciphering cell-type resolved cis-regulatory sequence lexicons with deep learning models of base-resolution chromatin accessibility profiles

To decipher the cis-regulatory sequence lexicon of TF binding sites in accessible cREs in each cell-type, we trained convolutional neural networks (called BPNet) to accurately map DNA sequence to base-resolution, pseudo-bulk chromatin accessibility profiles in 1 Kb windows around scATAC-seq peaks and in background regions (Avsec, Weilert, et al. 2021; Trevino et al. 2021) (**Figure 2a**). We used a 5-fold, chromosome hold-out cross-validation scheme to train, tune, and evaluate the predictive performance of the models (Trevino et al. 2021) (**Table S8-S9**). We obtained high and stable Spearman correlation between total predicted and observed Tn5 insertion coverage in test regions across all folds and cell types (**Figure 2b**, **Table S8**). The observed and predicted base-resolution distributions of Tn5 insertions (shapes of the profiles) in test peak regions were also concordant across folds and cell-types (**Figure 2b, Table S9**).

**Figure 2.**
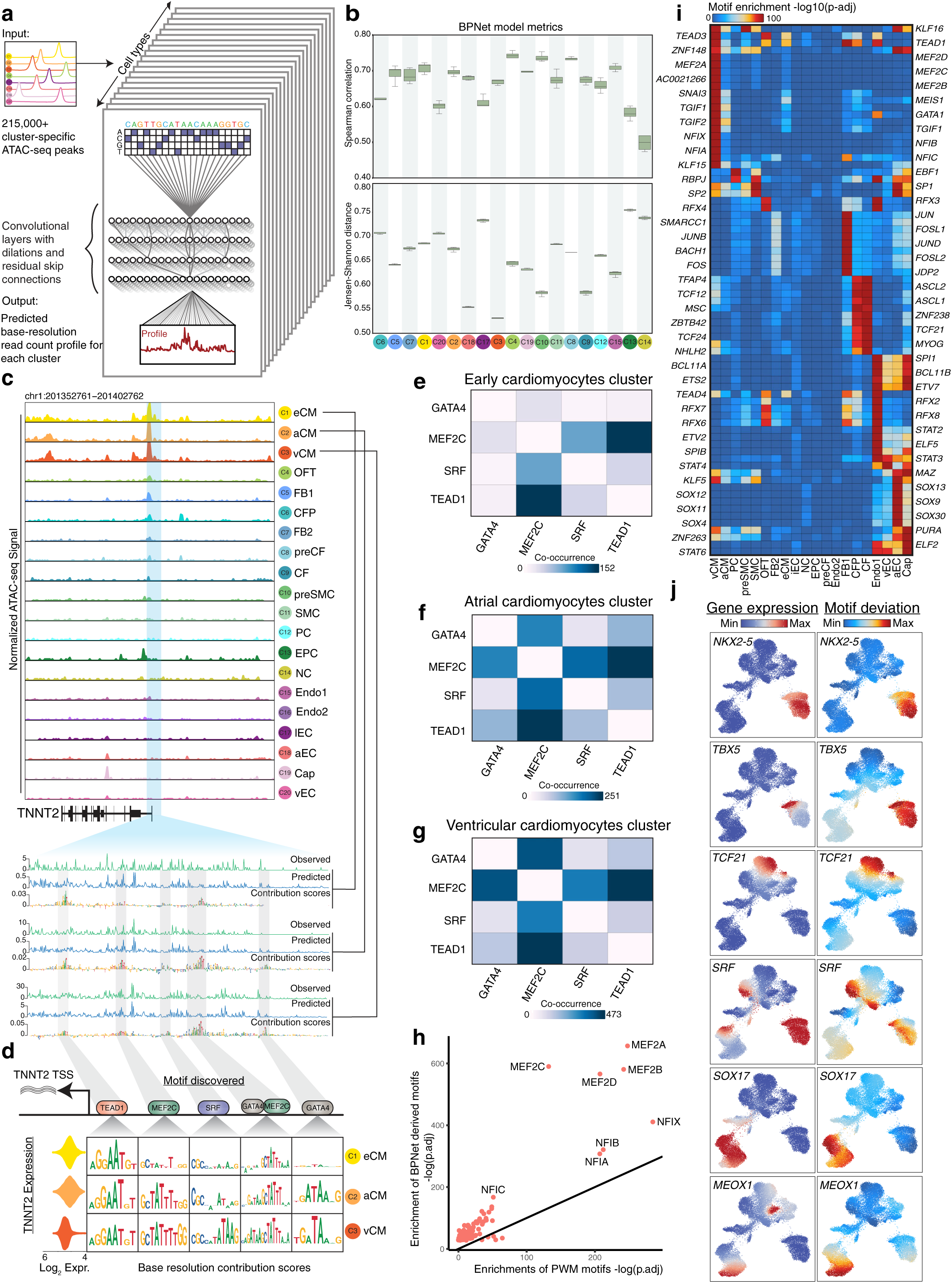
Cell-type resolved predictive transcription factor motif syntax derived from deep learning models of base-resolution scATAC-seq profiles. **(a)** Schematic of the convolutional neural network (BPNet) trained to simultaneously predict base-resolution probability distribution of reads and total read counts of cell-type resolved pseudobulk scATAC-seq profiles over each 1-kb accessible peak region from 2-kb underlying DNA sequences. **(b)** Performance evaluation of BPNet cluster-specific models, computed as the Spearman correlation between observed and predicted total counts (higher is better) across all peaks in each cluster (top) and mean Jenson-Shannon distance (lower is better) between the base-resolution observed and predicted profiles across all peaks in each cluster (bottom). Results are reported on test sets from a 5-fold cross-validation set up. **(c)** Top panel shows the genome tracks of aggregate pseudobulk scATAC-seq around the *TNNT2* locus for each of the cell-type clusters. The scale ranges from 0-0.34 in units of fold-enrichment relative to the total number of reads in TSSs per 10k. Bottom panel zooms into an accessible peak around the *TNNT2* transcription start site and shows the observed (Obs) base-resolution scATAC-seq read count profiles from the early (eCM), atrial (aCM) and ventricular cardiomyocytes (vCM) clusters, the predicted (Pred) profiles from the BPNet models of each of the three cell types and the corresponding DeepLIFT contribution score profiles (height of each base in the sequence is proportional to its contribution score). **(d)** Per-base DeepLIFT contribution scores of *TEAD1, MEF2C, SRF,* and *GATA4* motif locations in the *TNNT2* promoter from eCM, aCM and vCM (rows from top to bottom). Left-most column shows distribution of scRNA-seq expression (in units of log_2_(transcripts per 10K)) of TNNT2 across cells from each of the three clusters. **(e,f,g)** Pairwise motif co-occurrence counts for *TEAD1, MEF2C, SRF,* and *GATA4* motifs based on predicted active motifs across all accessible cREs in eCM, aCM and vCM respectively. **(h)** Comparison of statistical significance of overlap enrichment (-log *p*-value, Wilcoxon rank-sum test) of BPNet model-derived predictive motif instances (y-axis) vs. position weight matrix (PWM) based motif instances (x-axis) in vCM accessible peaks regions. Predictive motif instances show higher significance of enrichments. **(i)** Enrichments of BPNET model derived predictive motif instances of transcription factors (rows) in accessible peaks of different cell types (columns). **(j)** (left column) scRNA-seq gene expression (in units of log_2_(transcripts per 10K)) and (right column) scATAC-seq based ChromVAR motif deviation scores (in units of z-scores) for *NKX2-5* (scale for expr.: 0 to 6, motif dev.: -1 to 2)*, TBX5* (scale for expr.: 0 to 3, motif dev.: -1 to 2), *TCF21* (scale for expr.: 0 to 12, motif dev.: -1.5 to 3.5), *SRF* (scale for expr.: 0 to 0.4, motif dev.: -0.6 to 0.8), *SOX17* (scale for expr.: 0 to 1.8, motif dev.: -1 to 2) and *MEOX1* (scale for expr.: 0 to 1, motif dev.: -0.4 to 0.8) shown in the scATAC-seq UMAP representations of all cells.

Next, we interrogated BPNet models of each cell-type to infer predictive sequence features in each accessible cRE by using the DeepLIFT algorithm which derives the contribution of each nucleotide in the cRE sequence to the predicted accessibility (Shrikumar, Greenside, and Kundaje 2017; Lundberg and Lee 2017a). For example, DeepLIFT applied to the eCM BPNet model highlighted short, contiguous stretches of bases with high contribution scores, reminiscent of TF binding motifs, in the accessible promoter of *TNNT2*, a gene critical for sarcomere contractile function of the heart (Sehnert et al. 2002) (**Figure 2c**). Hence, we annotated predictive, active motif instances in all accessible cREs of each cell type by scanning their sequences with a non-redundant compendium of known TF sequence motifs (Weirauch et al. 2014) and restricting to instances with high DeepLIFT contribution scores or motif mutagenesis scores derived from each cell-type specific BPNet model. Although the sequence of a cRE is the same in all cell-types, its DeepLIFT contribution score profile can vary across cell types, reflecting cell-type specific prediction of motif activity by BPNet models of different cell types. For example, the *TNNT2* promoter is highly and equally accessible in all 3 types of cardiomyocytes (early (eCM), atrial (aCM) and ventricular (vCM)) and drives expression of *TNNT2* in all 3 cell types (**Figure 2c**). However, the DeepLIFT profiles derived from the eCM, aCM and vCM models for the same promoter sequence highlight distinct combinations of active TF motif instances predicted to regulate accessibility in the three cell types (**Figure 2c,d**). A *TEAD1* motif is predicted to regulate promoter accessibility in all three cell-types. A nearby *MEF2C* motif is predicted to be uniquely active in aCM and vCM, while another upstream *MEF2C* motif active in eCM is predicted to be part of a *GATA*-*MEF* composite motif that is specifically active in aCM and vCM. A *GATA* motif, further upstream, is predicted to be active specifically in aCM and vCM. An *SRF* motif is predicted to be active only in vCM. The higher density of predicted active motifs in the *TNNT2* promoter in aCM and vCM compared to eCM is concordant with the higher expression of *TNNT2* in the former two cell types (**Figure 2d**). This combinatorial, cell-type specific motif syntax of these 4 TFs at the *TNNT2* promoter is consistent with the genome-wide co-occurrence statistics of their active motifs across all cREs in eCM, aCM and vCM (**Figure 2e,f,g, Table S10**).

We expected the dynamic active motif instances with high contribution scores derived from cell-type specific BPNet models to be more representative of context-specific TF occupancy than static motif instances identified by classical position weight matrix (PWM) motif scanning methods that only use sequence match scores. Indeed, we found that most TFs, including those belonging to the *MEF2* family (adjusted *p*-value < 1e-500, Benjamini-Hochberg (BH) corrected hypergeometric test) and *NFI* family (adjusted *p*-value < 1e-150, BH corrected hypergeometric test) that are expected to be active in vCMs showed significantly stronger enrichment of active motif instances relative to PWM motif instances in differential, cell-type specific vCM peaks (**Figure 2h**).

Next, we estimated the enrichment of active motif instances of TFs in accessible cREs of each cell-type to identify the TF regulators of cell-type resolved chromatin accessibility landscapes (**Figure 2i, SFigure 5**). We found that *MEF2*, *TGIF1*, *NFI* motif families were highly enriched in vCMs and *TGIF* and *KLF* families in aCMs. The eCMs had similar TF motifs as the vCMs and aCMs, albeit with weaker enrichments, suggesting this cluster is the progenitor population for later cardiomyocyte subtypes. The CFPs and CFs had similar motif enrichment for *TCF21*/TCF, *MYOG, MSC*, with CF gaining enrichment for *TEAD* and *NFI* families and implicating a second set of TFs that become active during CF maturation. The other fibroblast-like clusters (FB1 and FB2) had lower *TCF21* enrichments than the cardiac fibroblast clusters, but stronger enrichment for *JUN*, *FOS* and *JDP* motif families. The OFT cells exhibited strong *RFX* and *TEAD* motif enrichments, while preSMC exhibited weaker enrichments for the *RFX* and *KLF* families and stronger enrichment for motifs associated with proliferation like *SP* and *RBPJ*. These enrichments became substantially stronger in the SMCs at PCW19, and with the gain of new TF enrichments such as the *MEF2* family, indicating a continuum of TF motif activity promoting the SMC cell fate trajectory. The PCW6 endocardial cells (Endo1) had stronger TF activity for *ETV* and *STAT* families and weaker enrichments for the *SOX* family. The capillary (Cap) cells, which are thought to derive from the endocardium, were strongly enriched for *SOX* family motifs. The aEC clusters, similar to capillaries, exhibited enrichments for *SOX*, *FOS* and *JUN* motifs and also retained endocardium TF motifs like *ELF* and *ETV*, while vEC had a motif landscape similar to the capillaries, with the addition of a few motifs, such as *STAT*. The cell type specificity of globally predictive TFs identified by the BPNet models was further corroborated by high concordance (**Table S11**) between TF activity scores (chromVAR (Schep et al. 2017)) and the expression of the TFs in the scRNA-seq data across developmental timepoints (**Figure 2j**). Our analyses thus provide a comprehensive resource of cell-type resolved TF lexicons and annotations of predictive TF sequence motifs in cRE landscapes of human fetal heart development.

### Inferring dynamic regulatory control across major cellular differentiation trajectories in human cardiogenesis

Next, we sought to identify major developmental trajectories involving cell state transitions across fetal heart development based on single-cell chromatin dynamics. We used the optimal transport algorithm (Schiebinger et al. 2019), previously developed to derive trajectories from single-cell gene expression data, to identify the most parsimonious transitions in global chromatin accessibility between cells from PCW6 to PCW19 of fetal heart development (**Figure 3a,b,c, SFigure 6, Table S12, Methods).**

**Figure 3.**
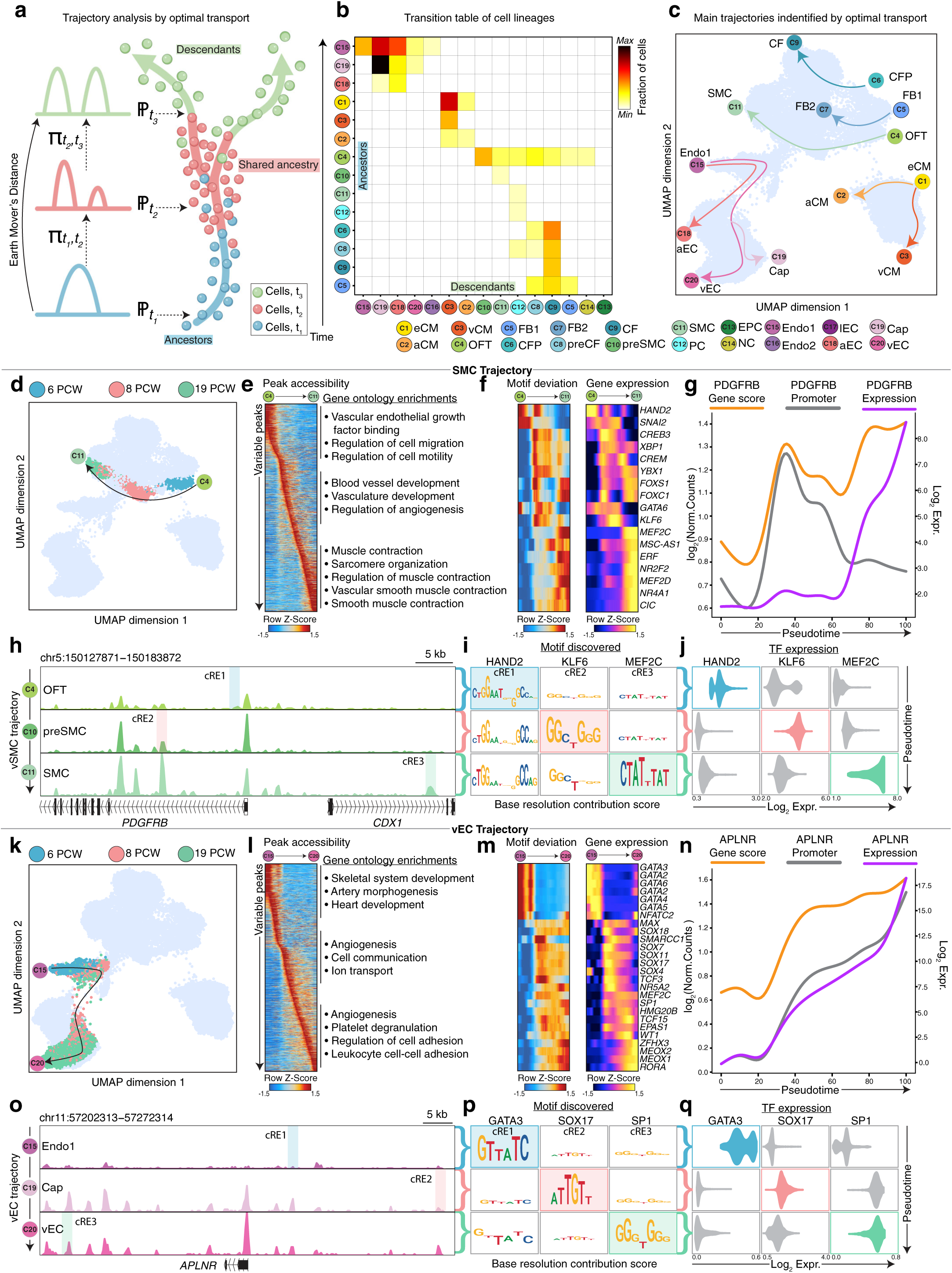
Identifying developmental trajectories in human fetal heart development. **(a)** Schematic of the optimal transport method used to determine trajectories of cell state transitions using scATAC-seq peaks of all the cell-types identified in Figure 1b. **(b)** Cell state transition table of cell lineages identified in the major trajectories obtained through optimal transport. Rows correspond to the parent cell-types and columns correspond to the derivative cell types. The heatmap is colored by the fraction of parent cells identified to be ancestors of the derivative cells. (scale for transition table.: 0.01 to 0.30) **(c)** UMAP of scATAC-seq cells highlighting the dominant trajectories identified using optimal transport. The cell-types correspond to those in Figure 3b. **(d)** UMAPs of scATAC-seq cells in the vascular smooth muscle cell (SMC) trajectory colored by the gestational sample time. **(e)** Heatmap of scATAC-seq signal (z-score of log_2_(reads per 10K)) of variable peaks identified in the SMC pseudotime trajectory. The gene ontology enrichments are calculated using the genes nearest to the variable peaks listed. **(f)** Heatmaps showing z-score of ChromVAR motif deviation scores (left) and gene expression log_2_(transcripts per 10K), also applicable for all gene expression values plotted in this figure) (right) of TFs with correlated variable activity in cells identified to be in the SMC trajectory, as ordered by pseudotime. **(g)** Gene expression, promoter chromatin accessibility log_2_(reads per 10K) +/- 500bp TSS and chromatin-derived gene accessibility score ((log_2_(reads per 10K), applicable for all gene activity values in this figure) dynamics of the *PDGFRB* gene across pseudotime. **(h)** Genome tracks of aggregate scATAC-seq data around the *PDGFRB* locus in OFT, preSMC and SMC clusters. cRE1, cRE2 and cRE3 are three representative cREs with dynamic motif activity further explored in (i) and (j). The ATAC signal range is 0-0.64 in units of fold-enrichment relative to the total number of reads in TSSs. **(i)** Per-base contribution scores of motifs of *HAND2*, *KLF6* and *MEF2C* in the 3 highlighted cREs in (h). Rows (top to bottom) are per-base contribution scores computed using BPNet models of OFT, preSMC, and SMC respectively. The columns (left to right) are the highlighted cREs from (h) that are active in OFT, preSMC, and SMC respectively. **(j)** Distribution of scRNA-seq gene expression of *HAND2*, *KLF6* and *MEF2C* TFs (columns) across cells from OFT, preSMC, and SMC clusters (rows). **(k)** UMAPs of scATAC-seq cells in the venous endothelial cell (vEC) trajectory colored by the gestational sample time. **(l)** Heatmap of z-scores of variable peaks identified in the vEC pseudotime trajectory. The gene ontology enrichments are calculated using the genes nearest to the variable peaks listed. **(m)** Heatmaps showing z-score motif activity (left) and normalized expression (right) of TFs with correlated variable activity in cells identified to be in the vEC trajectory, as ordered by pseudotime. **(n)** Gene expression, promoter chromatin accessibility and chromatin-derived gene accessibility score dynamics of the *APLNR* gene across pseudotime. **(o)** Genome tracks of aggregate scATAC-seq data around the *APLNR* locus in Endo1, Cap and vEC clusters. cRE1, cRE2 and cRE3 are three representative cREs with dynamic motif activity further explored in (p) and (q).The ATAC signal range is 0-0.52 in units of fold-enrichment relative to the total number of reads in TSSs per 10k. **(p)** Per-base contribution scores of motifs of *GATA3*, *SOX17* and *SP1* in the 3 highlighted cREs in (o). Rows (top to bottom) are per-base contribution scores computed using BPNet models of Endo1, Cap, and vEC respectively. The columns (left to right) are the highlighted cREs from (o) that are active in Endo1, Cap, and vEC respectively. **(q)** Distribution of scRNA-seq gene expression of *GATA3*, *SOX17* and *SP1* TFs (columns) across cells from Endo1, Cap, and vEC clusters (rows).

Within the endocardium lineage, the endocardium-like cell clusters (Endo1/2) were predicted to give rise to the capillary cells (Cap) which in turn were predicted to transition into the venous endothelium (vEC) cluster in PCW19. The aEC cluster was derived from Endo1/2 clusters as well as the PCW8 Cap cluster, suggesting that some terminal cell states can originate from different developmental origins (**Figure 3b**). We also identified cells that appeared to have already committed to their developmental fates based on their expression of lineage specific genes. For example, at PCW6, cells from the epicardial lineage (EPC, OFT, CFP and FB1) that expressed *TCF21* were predicted to transition into the cardiac fibroblasts at PCW8 (preCF) and PCW19 (CF) (**Figure 3b**). The OFT cluster which lacks *TCF21* expression was predicted to transition into SMC and PC clusters through the preSMC cluster. These observations are highly concordant with results from studies with lineage tracing in *TCF21* recombinase knock-in mice (Acharya et al. 2012). Finally, the FB1 cluster was predicted to transition into the FB2 cluster. For the myocardium cells, the eCM cluster was predicted to differentiate into vCM and aCM clusters. Overall, we characterized 8 dominant trajectories for all the major cell types at PCW19 (**Figure 3b,c, SFigure 7, SFigure 8**).

We then characterized genome-wide and locus-specific regulatory dynamics associated with cell state transitions across these trajectories. Below, we present representative case studies contrasting regulation of the development trajectories leading to SMC and vEC cell fates.

The SMC trajectory begins with the OFT cells at PCW6 that transition through an intermediate preSMC population in PCW8 to the SMCs at PCW19 (Davis et al. 2006) (**Figure 3d**). A continuous cascade of dynamically accessible cREs defines cell state transitions across the trajectory (**Figure 3e**). These dynamic cREs are proximal to genes enriched for temporally relevant vascular developmental processes including cell migration, angiogenesis, and muscle contraction at early, intermediate, and late time points, respectively (**Figure 3e**). Expression dynamics of several key lineage specifying TFs including *HAND2, SNAI2, KLF6* and *MEF2C* were strongly correlated with their chromatin-based motif activity (chromVAR deviation scores) across this trajectory (**Figure 3f**). Tracking the chromatin accessibility and gene expression of *PDGFRB*, one of the primary marker genes for the SMC population, we observed that initially, the promoter of *PDGFRB* accounts for the majority of accessibility at this locus while gene expression is low (**Figure 3g**) (Hellström et al. 1999; Levéen et al. 1994). The increase in expression of *PDGFRB* at later time points is associated with increased accessibility of putative intronic enhancers. We then used predictive motif instances derived from cell-type specific BPNet models to associate inferred TF binding dynamics at specific cREs in the *PDGFRB* locus with TF expression changes across the three timepoints (**Figure 3h,i**). BPNet models of OFT cells at the PCW6 time point revealed a predictive *HAND2* binding motif (**Figure 3i**) in a downstream putative enhancer (cRE1 in **Figure 3h**) that is highly accessible at this early time point. The predicted TF motif dynamics of *HAND2* at this enhancer was correlated with the expression dynamics of *HAND2,* which also peaks in PCW6 and decreases thereafter (**Figure 3j**). Another cRE (cRE2 in **Figure 3h**) proximal to the promoter of *PDGFRB*, which showed the highest accessibility in preSMC at the intermediate PCW8 time point, was predicted to be regulated by *KLF6* whose motif showed high contribution scores specifically in the preSMC model (**Figure 3i**) and whose expression also peaked in preSMCs (**Figure 3j**). A distal cRE upstream of *PDGFRB* (cRE3 in **Figure 3h**) with highest accessibility in SMC in PCW19 was predicted to be regulated by *MEF2C* whose motif was specifically predictive in SMC BPNet model (**Figure 3i**) and whose expression peaked in SMC (**Figure 3j**).

In contrast, the vEC differentiation trajectory captured cell state transitions from the Endo1/2 progenitor cells at PCW6 to vECs at PCW19 through the Cap cells in PCW8 (**Figure 3k**). Waves of TFs including *GATA2/3/4/6, NFATC2, SOX4, SOX17* and *MEOX1* with correlated expression and motif activity dynamics are predicted to regulate concordant cascades of dynamically accessible cREs targeting genes involved in different stages of angiogenesis (**Figure 3l,m**). We once again used cell-type specific BPNet models to decipher TFs that regulate dynamic cREs in the cis-regulatory domain of the *APLNR* gene, a primary marker of vECs (Sharma et al. 2017; Kang et al. 2013; Inui et al. 2006), which exhibited a coordinated and monotonic increase in gene expression, promoter accessibility and cumulative distal chromatin accessibility (gene accessibility scores) across the trajectory (**Figure 3n**). BPNet models trained on Endo1/2, Cap and vEC cells revealed *GATA3, SOX17* and *SP1* to specifically regulate three representative cREs in the *APLNR* locus with distinct temporal dynamics of chromatin accessibility based on cell-type specific predictive motif instances and concordant TF expression (**Figure 3o,p,q**). Our analysis framework thus provides a lens into the dynamic cis-regulatory code of developmental cellular trajectories in human cardiogenesis at unprecedented resolution.

### A systematic comparison of regulatory landscapes of *in vitro* differentiated cardiac cell types and their *in vivo* counterparts in human fetal heart development

While *in vivo* characterization of the regulatory programs of cell types and trajectories is critical for understanding cardiovascular development and disease pathogenesis, faithful cellular models that recapitulate human developmental processes are equally important, especially since they provide convenient, scalable platforms for experimental validation. Several human induced pluripotent stem cell (iPSC) based *in vitro* cellular models have been developed, including cardiomyocyte (i-CM), endothelial (i-EC), epicardial (i-EPC), cardiac fibroblast (i-CF), and smooth muscle (i-SMC) cells (Burridge et al. 2014; Lian et al. 2012; Cheung et al. 2012; H. Zhang et al. 2019). Our comprehensive, integrated single cell atlas of *in vivo* cardiac cell types from developing fetal hearts provides a unique opportunity to investigate the authenticity of these *in vitro* cellular models.

To address this question, we generated iPSC-derived i-CM, i-EC, i-EPC, i-CF, and i-SMC cells through directed differentiation employing established protocols (Burridge et al. 2014; Lian et al. 2012; Cheung et al. 2012; H. Zhang et al. 2019) (**Figure 4a**). There are three primary phases of *in vitro* differentiation of cardiac cell types: cardiac mesoderm induction from human iPSC, cardiac progenitor cell (i-CP) specification and proliferation, and i-CP differentiation directly to i-EC, i-EPC and primitive cardiomyocytes. i-EPCs were further differentiated into i-CFs and i-SMCs. Primitive cardiomyocytes (i-pCM) were differentiated into mature cardiomyocytes (i-CM) (Wamstad et al. 2012; J. Lee et al. 2018) (**Figure 4a**). We generated scATAC-seq data from all these *in vitro* differentiated iPSC lines at multiple time points, using the Chromium (10X Genomics) platform (**SFigure 9-10, Figure 4b, Table S13-S15**). Integration and clustering of cells from these scATAC-seq datasets broadly identified nine different cell types, including day 0 iPSC, day 2 mesodermal cells (i-Mes), day 5 i-CP, day 15 i-pCM, and day 30 i-CM, i-EPC, i-SMC, i-CF and i-EC. Once again, the scATAC-seq derived gene accessibility scores of marker genes were found to be highly specific for the relevant cell types, confirming our cell type annotations (Paik et al. 2018; Friedman et al. 2018; Churko et al. 2018) (**Figure 4c, SFigure 11, Table S15**). We also identified accessible peak regions as putative cREs and annotated TF motif instances in these cREs for each of the *in vitro* cell types (**SFigure 12**).

**Figure 4:**
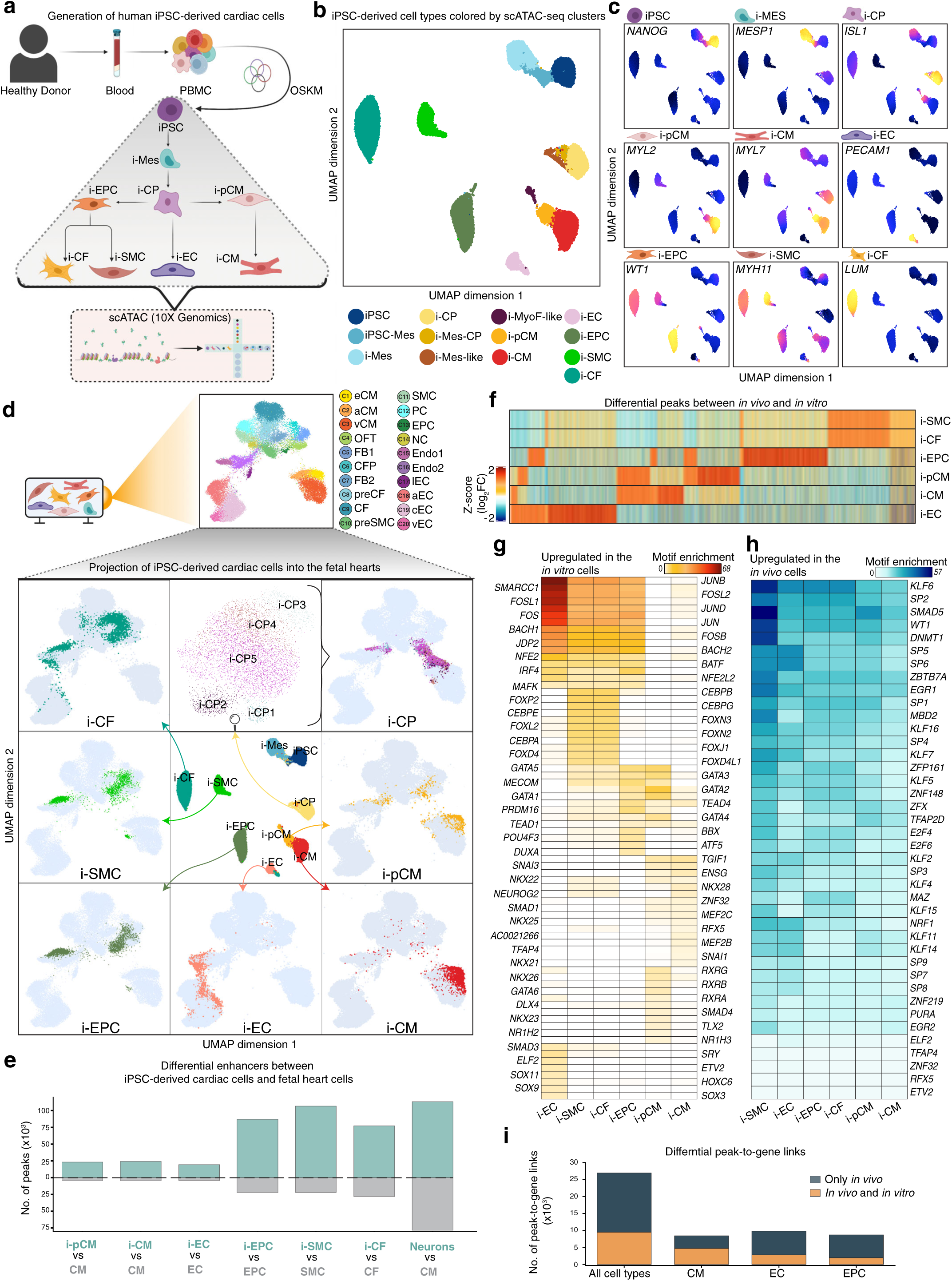
Characterization of *in vitro* iPSC-derived cardiac cell types. **(a)** Schematic for derivation of human iPS cells, followed by their differentiation to major cardiac cell types and genome-wide scATAC-seq profiling. **(b)** scATAC-seq UMAP of all *in vitro* iPSC-derived cells colored according to cell-types identified during differentiation (iPSC: induced pluripotent stem cells, i-PSC-Mes: partially differentiated mesoderm-like cells, i-Mes: cardiac mesoderm cells, i-CP: cardiac progenitors, i-Mes-CP: partially differentiated cardiac progenitor-like cells, i-Mes-End: partially differentiated endoderm-like cells, i-MyoF-like: Myofibroblast-like cells, i-pCM: Day 15 iPSC-derived primitive cardiomyocytes, i-CM: Day 30 iPSC-derived mature cardiomyocytes, i-EC: iPSC-derived endothelial cells, i-EPC: iPSC-derived epicardial cells, i-SMC: iPSC-derived smooth muscle cells & i-CF: iPSC-derived cardiac fibroblast cells). **(c)** Gene accessibility scores of marker genes *NANOG, MESP1, ISL1*, *MYL2, MYL7, PECAM1, WT1, MYH11* and *LUM* projected on the scATAC-seq fetal heart UMAP. **(d)** Projection of cells from scATAC-seq experiments profiling *in vitro* iPSC-derived cardiac cell types into the scATAC-seq fetal heart UMAP. Central panel in the 3×3 grid shows the scATAC-seq UMAP of all *in vitro* cardiac cell types. The other panels in the grid are projections of the i-CF (row 1, col 1), i-SMC (row 2, col 1), i-EPC (row 3, col 1), i-EC (row 3, col 2), i-CM (row 3, col 3) and i-pCM (row 2, col 3) cells into the scATAC-seq fetal heart UMAP. Panel in row 1, col 2 shows an scATAC-seq UMAP of 4 subclusters of cells from *in vitro* cardiac progenitors (i-CP1, i-CP2, i-CP3, i-CP4 and i-CP5) which are projected into the scATAC-seq fetal heart UMAP (row 1, col 3). **(e)** Comparison of number of significantly (log_2_ fold-change > 0.5, FDR < 0.01 using two-sided *t*-test) upregulated (in blue) and downregulated (in grey) scATAC-seq peaks in *in vitro* cardiac cell types relative to nearest *in vivo* fetal heart cell types. An analogous differential comparison between *in vivo* ventricular cardiomyocytes from fetal heart and *in vivo* glutamatergic neurons from fetal brain is shown as a reference (right-most bar). **(f)** Heatmap of *z*-score of differential scATAC-seq signal (log_2_ fold-change) of upregulated scATAC-seq peaks in *in vitro* cardiac types relative to nearest *in vivo* fetal heart cell types from panel (e). **(g,h)** Statistical significance (-log_10_(adjusted *p*-value), BH-adjusted hypergeometric test) of overlap enrichment of TF motifs in upregulated (g) and downregulated (h) scATAC-seq peaks in *in vitro* cardiac types relative to nearest *in vivo* fetal heart cell types from panel (e). **(i)** Number of peak-to-gene links in different *in vivo* fetal heart cell types that are also found (orange) in nearest *in vitro* cardiac cell types or exclusive (grey) to the *in vivo* cell types.

To evaluate the similarity between chromatin landscapes of the *in vitro* differentiated cell types and their *in vivo* counterparts, we first used the LSI method to project *in vitro* differentiated cells onto the scATAC-seq LSI subspace of all cells from the fetal heart samples (Granja et al. 2019) (**Figure 4d**). Majority of Day-15 primitive cardiomyocytes (i-pCM) projected into the PCW6 *in vivo* myocardium-derived cardiomyocytes (eCM). At day-30, *in vitro* cardiomyocytes (i-CM) projected primarily into the PCW8 *in vivo* ventricular cardiomyocytes (vCM) and *in vivo* early cardiomyocytes (eCM), while *in vitro* endothelial cells (i-EC) projected across the *in vivo* endocardial cells (Endo1, Endo2) and the PCW8 capillaries (Cap). In contrast, *in vitro* epicardium-derived cells, including i-EPC, i-SMC and i-CF, were distributed across epicardial cell types of the fetal heart without a strong correspondence to their specific *in vivo* counterparts (EPC, SMC and CF). The day-5 *in vitro* cardiac progenitors (i-CP) were found to consist of four sub clusters that projected across all three distinct lineages of the fetal heart, the myocardium, epicardium and endocardium, supporting the likely origin of all major differentiated *in vivo* cardiac cell types from a precursor state similar to i-CPs (**Figure 4d**).

To further corroborate the observed differences between *in vitro* cell types and their *in vivo* counterparts from the LSI co-projection, we identified scATAC-seq peaks with significant differential accessibility between matched *in vitro* and *in vivo* cell types (**Figure 4e**). As expected, *in vitro* i-pCMs, i-CMs, and i-ECs had the least number of differential peaks relative to their matched *in vivo* cell types (**Figure 4e**). Consistent with the co-projection analysis, comparison of matched *in vitro* epicardial cell types (i-EPC, i-SMC and i-CF) and their *in vivo* counterparts revealed more differential peaks relative to corresponding comparisons of cardiomyocytes and endothelial cells. To calibrate the magnitude of these differences, we also estimated differential peaks between two distant *in vivo* cell types, namely vCMs and excitatory neurons (Trevino et al. 2021). Reassuringly, the differences between *in vitro* and *in vivo* epicardial cells were substantially smaller than differences between vCMs and neurons (**Figure 4e**).

Next, to study the regulatory basis of the differences in chromatin landscapes between *in vitro* and *in vivo* cardiac cell types, we identified TF motifs enriched in the differentially accessible scATAC-seq peaks (**Figure 4f**). *AP-1* (*JUN-FOS, JDP2*) motifs were strongly enriched in peaks upregulated in most *in vitro* cell types, except cardiomyocytes (**Figure 4g**). In contrast, downregulated peaks in *in vitro* cell types were most enriched for *SP, KLF* and *WT1* motifs (**Figure 4h**). Differentially upregulated peaks in *in vitro* cardiomyocytes (i-pCM, i-CM) were enriched for motifs of classical cardiac TFs including *MEF2* and *NKX*, consistent with their role in cardiomyocyte differentiation (Vincentz et al. 2008). Motifs of *FOX* and *CEBP* TF families, which are involved in epithelial-to-mesenchymal transition (EMT), were enriched in peaks upregulated in *in vitro* epicardium-derived cell types compared to their post-EMT *in vivo* counterparts (Quijada, Trembley, and Small 2020; von Gise and Pu 2012; Risebro, Vieira, and Riley 2015; Lamouille, Xu, and Derynck 2014), suggesting that the *in vitro* epicardial cells may not represent a terminally differentiated state.

Finally, to understand the differences in regulatory enhancer-gene networks between matched *in vitro* and *in vivo* cardiac types, we identified putative enhancer-gene links across *in vivo* cell cardiac types based on correlation of *in vivo* scATAC-seq signal at distal CREs with *in vivo* scRNA-seq expression of neighboring genes. We integrated our scATAC-seq data from the *in vitro* differentiated cell types with previously published scRNA-seq data from similar *in vitro* differentiation experiments (Friedman et al. 2018) (**SFigure 13, Table S16**) in order to derive analogous enhancer-gene links across the *in vitro* cell types (Granja et al. 2019) (**SFigure 14, Table S17, S18**). *In vitro* and *in vivo* cardiomyocytes shared the highest proportion of enhancer-gene links, followed by endothelial cells, consistent with the strong concordance of their chromatin landscapes (**Figure 4i**). These comparative analyses collectively suggest that *in vitro* cardiomyocytes and endothelial cells may serve as representative model systems for their *in vivo* counterparts in fetal heart development.

### Prioritizing putative causal non-coding genetic variants, TFs, target genes and cell types in cardiovascular disorders and congenital heart diseases

Common genetic variants associated with complex traits and diseases have been previously found to be strongly enriched in cREs of disease-relevant cell types and tissues (Maurano et al. 2012; Schaub et al. 2012; Finucane et al. 2015; Claussnitzer et al. 2020). Hence, we used stratified linkage disequilibrium (LD) score regression to estimate the proportion of heritability from GWAS summary statistics of cardiovascular and other control diseases and traits that can be attributed to the accessible cRE landscape of each *in vivo* cardiac cell type in our atlas (Bulik-Sullivan et al. 2015; Nielsen et al. 2018) (**SFigure 15a, Table S19,S20**). Variants associated with atrial fibrillation were significantly enriched (enrichment = 33.215, Bonferroni adjusted *p*-value = 2.166e-6) in accessible cREs of all three types of *in vivo* cardiomyocytes with the strongest enrichment in atrial cardiomyocytes (aCM). aCM cREs were also significantly enriched (enrichment = 15.163, Bonferroni adjusted *p*-value = 0.01798) for GWAS loci associated with heart failure phenotypes. Coronary artery disease GWAS loci were significantly enriched (Bonferroni adjusted *p*-value < 0.05) in cREs of multiple cell types (vCM, preCF, CF, SMC, PC, Endo1/2, Cap, vEC and aEC) with the highest enrichment (enrichment = 20.753, *p*-value = 8.987e-4) in arterial endothelial cells (aEC). GWAS signal from control phenotypes such as inflammatory bowel disease was not enriched in the regulatory landscape of any of our cell types, indicating specificity of these cell types for cardiac traits and disease.

Next, we investigated the utility of our regulatory atlas of cardiogenesis to decipher causal single-nucleotide, *de novo*, non-coding mutations in congenital heart disease (CHD) patients. We compiled a set of 54,126 *de novo*, non-coding point mutations from 763 CHD patients from the Pediatric Cardiac Genomics Consortium (Richter et al. 2020) (PCGC) (**Table S21**) and a control set of 110,055 *de novo*, non-coding point mutations from healthy controls from the Simons Simplex Collection (n=1902 trios) (**Table S22**). We tested the accessible cRE landscapes of each of the *in vivo* fetal heart cell types for enrichment of case versus control mutations. Surprisingly, only two cell types, Capillaries (Cap) and Endocardium (Endo1), were barely enriched (odds-ratio (OR) = 1.023 and 1.021 respectively) and all other cell types lacked enrichment (OR < 1) (**SFigure 15b**). These results suggest that overlapping mutations with cell-type resolved cREs is not sufficient to prioritize potentially causal CHD mutations.

Previous studies have successfully nominated functional non-coding disease variants by leveraging predictive sequence models of chromatin state in relevant cell types and tissues (Avsec, Agarwal, et al. 2021; David R. Kelley, Snoek, and Rinn 2016; D. R. Kelley et al. 2018; D. R. Kelley 2020; J. Zhou and Troyanskaya 2015; Richter et al. 2020; J. Zhou et al. 2018; Trevino et al. 2021). Hence, we hypothesized that cell-type specific mutation impact scores inferred from BPNet models of chromatin accessibility in fetal heart cell types might improve prioritization of causal *de novo*, non-coding CHD mutations and their cell types of origin. For each cell type, we used the corresponding BPNet models to estimate mutation impact scores of candidate case and control mutations in accessible cREs as the log_2_ fold-change in the cumulative predicted scATAC-seq profile probabilities for both alleles over a 100 bp window centered at each mutation (**Figure 5a**). We observed striking variation of enrichment of mutations with high predicted mutation impact scores in cases versus controls across the cell types (**Figure 5b**). Mutations prioritized in several cell types showed weak to moderate enrichments including NC (OR = 1.016), lEC (OR = 1.033), EPC (OR = 1.042), Endo1 (OR = 1.106), vEC (OR = 1.092), vCM (OR = 1.119), Cap (OR = 1.205), OFT (OR = 1.22) and preSMC (OR = 1.307) (**Figure 5b, Table S23,24**). The highest enrichment (Cases n = 47; Control n = 56; OR = 1.707; *p-*value = 0.008, Fisher’s Exact test) was obtained for mutations prioritized in arterial endothelial cells (aECs) (**Figure 5b,c,Table S23,24**), which is consistent with the contribution of this endothelial cellular lineage to multiple cardiac structures. These patterns of cell-type specific enrichment were robust to different measures of mutation impact scores and thresholds for defining high impact mutations (**SFigure 15c-f**). These results suggest that mutation impact scores from BPNet models trained on cell-type resolved chromatin accessibility profiles are able to prioritize putative causal CHD variants in relevant cell types.

**Figure 5:**
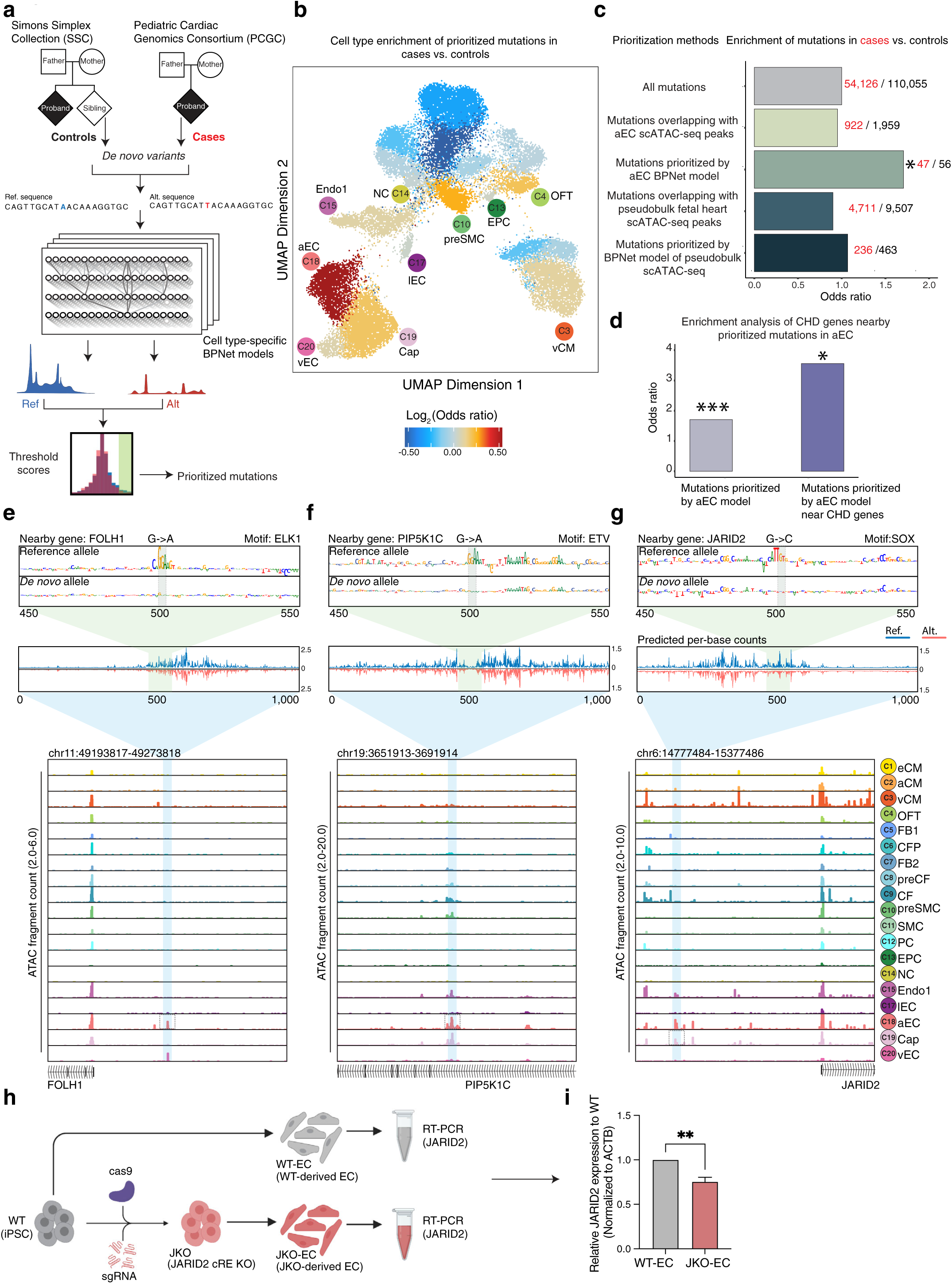
Prioritizing non-coding CHD mutations using deep learning models of scATAC-seq profiles from fetal heart cell types. **(a)** Schematic of mutation prioritization pipeline that uses cell-type specific BPNet models to predict scATAC-seq profiles for sequences containing both alleles of candidate mutations and derive a mutation impact score. The models were used to score CHD-associated *de-novo* non-coding mutations from the PCGC cohort and mutations from healthy controls from the SSC cohort. **(b)** Enrichment (log2(OR), Fisher’s Exact Test) of prioritized mutations from each cell-type specific BPNet model in CHD cases vs. controls plotted on the scATAC-seq UMAP of all fetal heart cells. **(c)** Enrichment of mutations in CHD cases vs. controls prioritized using different methods. Mutations prioritized by BPNet models trained on arterial endothelial (aEC) scATAC-seq profiles are significantly enriched in cases vs. controls (OR = 1.707, *p*-value = 0.008, Fisher’s exact test). **(d)** Enrichment of mutations prioritized by aEC BPNet model in cases vs. controls (grey bar) compared to enrichment of mutations prioritized by aEC BPNet model proximal to CHD associated genes (blue bar) in cases vs. controls. **(e,f,g)** Case studies of three prioritized *de-novo* CHD mutations in endothelial cREs in the *FOLH1* (e), *PIP5K1C* (f) and *JARID2* (g) gene loci respectively. Top-most panel shows contribution scores derived from cell-type specific BPNet models (aEC for (e,f) and Cap for (g)) of each nucleotide in a 100 bp sequence window containing each allele of the mutation. The changes in contribution scores highlight disruption of active TF motifs (ELK/ETV motifs for (e,f) and SOX motif for (g)). The panel below shows corresponding predicted base-resolution scATAC-seq count profiles in a 1 Kb window containing reference (blue) and alternate (red) allele of the mutation (the red tracks for the alternate alleles are inverted along the x-axis). These tracks highlight local disruption of predicted scATAC-seq profiles by the mutations. The last panel shows observed cell-type resolved pseudobulk scATAC-seq coverage profiles for all cell types at each locus. Scale of tracks is 2.0-6.0 (*FOLH1*), 2.0-20 (*PIP5K1C*) and 2.0-10.0 (*JARID2*) in units of Tn5 insertion counts observed in each cell type. **(h)** Schematic of *in vitro* differentiation of iPSCs to EC lineage, and comparison of *JARID2* expression in iPSC-derived EC with and without CRISPR-Cas9 deletion of cRE containing prioritized CHD mutation shown in (g). **(i)** CRISPR-Cas9 deletion of cRE containing the prioritized CHD mutation from (g) shows significant decrease (**p < 0.001, two-sided t test) in expression of *JARID2* gene expression in knockout vs. wild-type iPSC-derived ECs.

In contrast, mutation impact scores derived from BPNet models trained on pseudobulk scATAC-seq profiles agglomerated over all fetal heart cell types did not enrich for CHD mutations (OR = 1.01), indicating that cell-type specificity of mutation impact scores is critical for prioritizing *de novo* CHD mutations (**Figure 5c**). To understand the benefits of quantitative, base-resolution profile BPNet models over conventional, peak-resolution, binary classification models for mutation impact prediction (Richter et al. 2020; J. Zhou and Troyanskaya 2015; David R. Kelley, Snoek, and Rinn 2016), we also estimated mutation impact scores from a neural network with the same base architecture as BPNet trained on binary, peak-resolution accessibility in the aEC cell type. Mutation impact scores from the binary aEC model showed weaker enrichments and statistical significance (OR = 1.448, p-value = 0.056) compared to those obtained from the base-resolution, profile BPNet aEC model (OR = 1.707; *p-*value = 0.008) (**SFigure 15g-i, 16a**). Mutation impact scores from our BPNet profile models also substantially outperformed scores derived from a previous peak-resolution, binary neural network model called HeartENN (Richter et al. 2020) which was trained on a large compendium of bulk chromatin data and used to score CHD mutations (**SFigure 16a**). These results suggest that cell-type resolved, chromatin accessibility profiles of the developing fetal heart as well as mutation impact scores inferred from base-resolution profile BPNet models trained on these data are both critical for enhancing prioritization of non-coding CHD mutations and affected cell types.

We further examined whether high impact mutations prioritized by BPNet in aECs occurred near genes previously associated with CHD based on genetic studies in human cohorts or mouse models obtained from (Richter et al. 2020) (744 total CHD-associated genes). We observed a 3-fold enrichment (*p*-value = 0.0486, Fisher’s Exact test) of predicted high-impact aEC mutations proximal to previously implicated CHD genes in cases (n = 7 of 47) compared to controls (n = 4 of 56) (**Figure 5d**).

Next, we performed deeper investigations of the causal chain of TF binding sites, cREs and target genes potentially affected by a subset of high-impact CHD mutations prioritized in aECs that are in close proximity (< 200 bp) to summits of high coverage aEC scATAC-seq peaks (**Table S25**). We used the active motif annotations derived from the cell-type specific BPNet models and the corresponding allele-specific base-resolution contribution scores of cRE sequences harboring these mutations to infer potentially disrupted TF binding sites (**Figure 5e,f,g**). A prioritized G-to-A *de-novo* mutation was predicted to ablate an *ELK/ETV* TF motif in a cRE that is exclusively accessible in endothelial cells (aEC, Cap, vEC and lEC) and ∼25 Kb upstream of a folate hydrolase gene *FOLH1*. *FOLH1* is expressed in endothelial cells (**SFigure 16b**) and has been associated with loss of normal structural endothelial cell integrity (Eppig et al. 2015; Conway et al. 2006) (**Figure 5e**). Another G-to-A mutation was predicted to disrupt an *ELK/ETV* TF motif in an endothelial cRE in the intron of the *PIP5K1C* gene, an important developmental TF strongly expressed in endothelial cells (**SFigure 16c**) and implicated in cardinal vein and right ventricular development and CHD (Eppig et al. 2015; Y. Wang et al. 2007; Richter et al. 2020) (**Figure 5f**). Interestingly, several other prioritized mutations were also predicted to disrupt *ELK/ETV* binding sites in accessible aEC cREs proximal to the *MGAT1*, *TIMP3, TBX3* and *NEK3* genes (**Table S25**), all of which have been previously associated with CHD or cardiovascular defects in human genetic studies or mouse models (Richter et al. 2020; Yuan Zhang et al. 2020; Eppig et al. 2015; Fedak et al. 2004; Kawamoto et al. 2006; Mesbah et al. 2008). We also found a G-to-C mutation in an accessible cRE distal to the *JARID2* gene predicted to disrupt a *SOX* TF motif in aEC and Cap cells (**Figure 5g**). JARID2 is an important endothelial TF (**SFigure 16d**) during early heart development, and coding mutations in *JARID2* have been implicated in CHD by previous studies, especially for tetralogy of Fallot (Mysliwiec, Bresnick, and Lee 2011; Cho et al. 2018; Y. Lee et al. 2000; Barth et al. 2010). In order to experimentally verify the functional impact of the cRE overlapping this prioritized mutation on *JARID2* gene expression in endothelial cells, we used CRISPR/Cas9 to delete 352 bp around the mutation in iPSCs, selected single clones with bi-allelic deletions of the targeted locus, differentiated these clones into endothelial cells and measured expression of *JARID2* (**Figure 5h, SFigure 17**). We observed a significant decrease (1.3-fold, *p*-value < 0.001, two-sided t test) in *JARID2* expression (**Figure 5i**) in edited iPSC-derived ECs compared to the isogenic controls. This experiment demonstrates that this cRE harboring the prioritized CHD mutation modulates the expression of its putative causal gene in the nominated cell type. In principle, the change in expression of this and other critical genes could cause significant downstream transcriptional changes that in turn affect cellular phenotypes leading to CHD. Our analysis framework thus enabled us to prioritize putative causal, *de-novo* non-coding mutations, their putative target TF binding sites and genes as well as the relevant cell types within the developmental window that are likely to be affected by these mutations potentially causing CHD.

## Discussion

In this study, we present a unique resource elucidating regulatory dynamics of human cardiogenesis at single cell resolution. By generating scATAC-seq experiments in fetal hearts at early and mid-gestational developmental time points and integrating these data with complementary scRNA-seq experiments from previous studies, we reveal the coordinated landscapes of dynamic cREs and genes that define major cell types, lineages and differentiation trajectories in the developing human heart. The single cell nature of our atlas allows identification of even subtle regulatory heterogeneity in progenitor populations that might represent lineage committed progenitors. A salient example of this is the observation of two distinct post EMT populations one of which has high TCF21 gene activity, and the other with low activity. Our trajectory analysis predicts this high activity subtype to be the progenitor of the fibroblast lineage, while the low TCF21 population is predicted to be progenitors of the vascular smooth muscle lineage (**SFigure 3**). The identification of these epigenetically distinct progenitor populations may represent lineage specification events that are especially important for understanding disease causality.

By training and interpreting deep learning models that accurately predict base-resolution profiles of cell-type resolved chromatin accessibility profiles from the underlying DNA sequence, we are able to decipher the cell-type specific sequence syntax of active TF binding sites in all cREs of major cardiac cell types at unprecedented resolution. These cell-type specific models of regulatory DNA do not simply identify all enriched TF motif instances in accessible chromatin, but rather learn and enable *ab initio* discovery of active TF motif instances and higher-order motif syntax predictive of the shape and strength of chromatin accessibility profiles at individual elements in each cell type. By coupling these dynamic TF motif activity maps with TF expression across the cell types, we define putative trans-factors that bind to TF motif syntax encoded in specific cREs and orchestrate dynamic gene expression programs that define differentiation trajectories of the major cardiac cell types. We identify diverse previously-validated TFs in mice that are important for cell fate determination of the terminally differentiated cell types. For example, we identified Sox17 to be a TF with predicted dynamic binding in the late capillary (**SFigure 8c**) and mid venous (**Figure 3m**) differentiation trajectory in open chromatin peaks near *APLNR* (**Figure 3p**). Consistent with these findings, *SOX17* knockout in mice has been shown to retard differentiation of endocardial cells due to downregulation of the NOTCH signaling pathway and promote defective heart development (Saba et al. 2019).

These observations demonstrate the temporal cell-restricted expression of the *SOX17* developmental TF, and implicate the NOTCH regulated APELIN-APLNR pathway (Helker et al. 2020) in *SOX17* directed endocardial development. Likewise, *TCF21* has been shown to be a key developmental TF regulating the formation of cardiac fibroblast cells during cardiogenesis (Acharya et al. 2012), with a putative role in regulating chromatin accessibility at enhancers and enabling the binding of other related TFs along the complex trajectory of precursor cells in the early to middle stages of cardiac fibroblast differentiation (**SFigure 7h-k**). Consistent with these findings, we observe *TCF21* to be an early to mid-regulator of the cardiac fibroblast trajectory. We also observe molecular signatures of activity for *MEF2C (Desjardins and Naya 2016)* in ventricular cardiomyocytes, *GATA4* in atrial cardiomyocytes (Misra et al. 2012) and *HAND2* in vascular smooth muscle cells (Barnes et al. 2011). Apart from revealing the precise timing of activity at cREs for these known master regulators, we also nominate putative novel regulatory TFs. For example, we observe *SOX18* expression and chromatin activity in the mid to late temporal regulation of arterial endothelial cells. This activity pattern is consistent with other data implicating this factor, along with *SOX17*, in regulating vascular endothelium development in mouse retina (Y. Zhou et al. 2015) (**SFigure 8g**) and controlling the expression of *MEOX2 (Douville et al. 2011)* and *CLDN5* -- downstream master regulators of arterial development (Fontijn et al. 2008) (**SFigure 8c**). We also identify other TFs that exhibit strong chromatin activity changes along developmental lineage trajectories (**Figure 3f,m, SFigure 7, SFigure 8**), implicating these factors as potentially important for lineage specification.

*In vitro* differentiated cardiac cell types are commonly used as convenient, scalable experimental models of *in vivo* cardiac cell types. However, the molecular authenticity of these *in vitro* cellular models as surrogates for studying *in vivo* cardiac development has yet to be verified. By augmenting our cell-type resolved regulatory atlas of fetal heart development with scATAC-seq profiling of diverse *in vitro* differentiated cardiac cell types, we were able to perform a systematic analysis of the differences of regulatory landscapes between these *in vitro* differentiated cardiac cell types and their *in vivo* counterparts. While the best *in vitro*-derived cellular proxies were cardiomyocytes and endothelial cells, we observed that epicardial derived lineages, including cardiac fibroblasts and SMC, exhibited much stronger EMT related TF activities than their *in vivo* counterparts (**Figure 4g**). The EMT program enables cells to acquire a non-terminally differentiated phenotype (Kalluri and Weinberg 2009), and in our data appears to drive substantial large-scale epigenetic differences in the *in vitro* derived epicardial lineage cells compared to their *in vivo* nearest neighbors. We hypothesize that inhibiting these EMT related TFs, for example by inhibiting the TGFβ pathway (Kalluri and Weinberg 2009), might improve the epicardial lineage differentiation protocols. Also, we believe that this single cell molecular “benchmarking” against *in vivo* derived data will become a useful computational tool in optimizing future variations of *in vitro* differentiation protocols.

Finally, our study also showcases the power of integrating single cell regulatory maps of cardiac development with cell-type specific deep learning models of regulatory DNA sequence to provide new insights into the genetic, molecular and cellular basis of congenital heart disorders (CHD). By using the deep learning models as *in-silico* oracles, we predict the impact of candidate non-coding variants and *de novo* mutations on cell-type specific chromatin accessibility profiles and infer the active TF binding sites disrupted by high impact mutations. We not only prioritize likely causal, high impact, *de novo* non-coding mutations in CHD patients but also identify likely causal cell types whose cREs are strongly enriched for prioritized CHD mutations. Crucially, we show that overlap of mutations with cell-type resolved cRE maps of fetal heart cell types is not sufficient to enrich CHD mutations, unless augmented by mutation impact scores from our cell-type specific deep learning models. Moreover, the cell-type specificity of our models as well as their ability to make base-resolution quantitative predictions of chromatin accessibility are both critical to the success of this approach. Mutation impact scores derived from models trained on a pseudo bulk aggregation of all our fetal heart cell types (a reasonable proxy for bulk tissue open chromatin data sets) or other bulk chromatin profiling datasets of cardiac tissue were unable to enrich for *de novo* mutations in CHD cases relative to healthy controls (**Figure 5d**). This observation strongly suggests that disruption of regulatory networks by *de novo* mutations is highly cell type specific, and inclusion of unaffected cell types leads to poorer prioritization of the predicted deleterious mutations. Our CRISPR/Cas9 validation experiments in *in vitro* differentiated endothelial cells ablating an endothelial lineage specific enhancer harboring a predicted high impact *de novo* CHD mutation provides evidence that prioritized mutations likely impact bona-fide enhancers. Importantly, our predictive framework for prioritizing CHD mutations has identified the arterial endothelial cell as harboring the greatest enrichment for CHD risk, suggesting that these cells play an important role in the structural specification of the heart. While previous studies have identified some putative arterial CHD causal variants and genes, they have not been able to assign the putative causal mutations and genes to specific cell lineages or developmental trajectories (Richter et al. 2020). Arterial endothelial cells provide the oxygen and nutrients necessary for heart morphogenesis, and thus disease related pathways may reflect a circulatory function, but paracrine signaling to other developmental cell types is also a possibility.

However, our work does have some limitations and caveats. First, while most developmental trajectories exhibited no substantial “gaps” in cell density, obtaining samples both earlier and later in development might allow us to more fully populate the extremes of these trajectories, extending our understanding of these developmental trajectories. Second, our analysis of regulatory landscapes has largely focused on activators, as repressors are both more challenging to nominate using correlation-based analysis, and may have activity that is less correlated either with increasing or decreasing accessibility. Third, we restrict our prioritization of *de novo* CHD mutations to those that fall in the immediate vicinity of observed scATAC-seq peaks in our fetal heart atlas and are likely to disrupt and decrease accessibility. While this strategy reduces the likelihood of false positives, it does bias our prioritization against mutations that might result in gain of accessibility. The reduced sensitivity of peak identification from scATAC-seq profiles in some rare cell types (e.g. neural crest cells) with sparse coverage may also result in a greater false negative rate and reduced enrichments for these cell types. We also hope to expand our analyses in the future to other types of mutations beyond point mutations. Finally, while we have directly validated the impact of one candidate enhancer harboring a specific *de novo* CHD mutation on expression of its predicted target gene, more extensive computational and experimental validation of the gene expression impact of prioritized mutations would further delineate the “hit rate” of our models, and also begin to characterize the distribution of the magnitude of gene expression effects these high-impact mutations might have.

In summary, we present an integrative framework that couples scATAC-seq and scRNA-seq experiments in developing organ systems with cell-type resolved deep learning models of regulatory DNA to dissect dynamic, cis and trans regulatory programs at unprecented resolution, and to decipher the genetic, regulatory and cellular drivers of related developmental disorders. We have previously demonstrated the utility of this framework to dissect regulation of human corticogenesis and prioritize *de novo* non-coding mutations in autism (Trevino et al. 2021). Our current study showcases the generalizability of the same framework to study regulation of human cardiogenesis and successfully prioritize causal *de novo* non-coding drivers of CHD. In the future, given the availability of multi-omic measurements from the same nuclei (Ma et al. 2020), we expect further improvements to these predictive models enabling quantitative predictions of the effects of non-coding mutation on cell-type specific chromatin state and gene expression (Avsec, Agarwal, et al. 2021; J. Zhou et al. 2018; Karbalayghareh, Sahin, and Leslie 2021). Such advances will further illuminate the mechanisms by which *de novo* non-coding mutations might manifest CHD.

## Acknowledgements

We thank members of the Kundaje, Greenleaf, Quertermous, Wang, Wu, Engreitz and Karakikes labs for discussion and advice, especially J. Granja and G. Marinov. We also thank Hilary Finucance for sharing the data used in linkage disequilibrium score analysis. All schematics were created with BioRender. Sequencing of scATAC-seq libraries was performed by the Stanford Functional Genomics Facility (supported by NIH grants S10OD025212 and 1S10OD021763).

## Author Contributions

M.A., L.S., I.K., K.C.W. and A.K. conceived the project. L.S., M.A., T.Q, W.J.G and A.K. generated figures and wrote the manuscript with input from authors. M.A. designed and performed all experimental data generation for the manuscript with inputs from L.S., M.C., K.D.W, M.G., I.K., K.C.W., T.Q., A.K. and W.J.G.. L.S designed and performed all analyses for the manuscript with inputs from M.A, A.Ban., S.K., S.N., A.S., A.V., N.V., A.Bal, J.E., K.F., T.Q, W.J.G & A.K..

## Data availability

Aligned fragment files from single cell chromatin assays are deposited in the Gene Expression Omnibus database with the SuperSeries reference number GSE181346.

The cell by gene accessibility scores matrices along with cluster 5’ insertion bigWig tracks for the human heart samples are deposited to UCSC cell browser portal under reference url : https://cardiogenesis-atac.cells.ucsc.edu to enable visualization of cell markers and genes for the broader community.

Code used for single cell analysis, training BPNet models and reproducing results for all figures can be found at: https://github.com/kundajelab/Cardiogenesis_Repo.

Interactive HiGlass browser sessions with cell-type resolved tracks for measured base-resolution scATAC-seq coverage profiles and predicted base-resolution scATAC-seq coverage profiles from BPNet models as well as model-derived nucleotide-resolution contribution scores in peak regions can be found at: https://resgen.io/kundaje-lab/sundaram-2022/views/cardiogenesis (Please press the Alt key or Option key + mouse scroll to scroll down or up through the tracks).

## Supplementary information

Methods

Figure Legends S1–S17

List of Supplementary Tables

## Experimental methods

### Patient recruitment

Human subjects were enrolled in the study with informed consent approved by the Stanford Institutional Review Board and Stem Cell Research Oversight Committee. Human fetal heart tissues (day-42, day-56, and day-133 post conception) were obtained from de-identified aborted fetuses in collaboration with the Stanford Family Planning Research Team, Department of Obstetrics and Gynecology, Division of Family Planning Services and Research, Stanford University School of Medicine. Human iPSCs were obtained from the Stanford CVI iPSC Biobank.

### Generation and culture of human-induced pluripotent stem cells

Peripheral blood mononuclear cells (PBMCs) were reprogrammed to hiPSCs using the CytoTune™-iPS 2.0 Sendai Reprogramming Kit (Thermo Fisher Scientific) according to the manufacturer’s instructions with modifications as previously described (Feyen et al. 2021). Stem cell-like colonies were manually picked about two weeks post-transduction and expanded in E8 stem cell media (Life Technologies). All iPSCs used for the subsequent studies were within passages 22 to 30. The genome integrity was assessed by a single nucleotide polymorphism-based karyotyping assay (Illumina, HumanOmniExpress-24 v1.1). The iPSCs were maintained in a defined E8 medium (Life Technologies) on cell culture plates coated with ESC-qualified Matrigel (BD Biosciences) in a hypoxic environment (8% O_2_, 5% CO_2_) at 37°C. For routine passaging, iPSCs were dissociated with Gentle Cell Dissociation Reagent (StemCell Technologies) and cultured on with E8 medium supplemented with 5 μM Y-27632 (SelleckChem). The iPSCs were tested to be mycoplasma negative using the Mycoalert Mycoplasma testing kits (LT07-318, Lonza).

### Cardiomyocyte differentiation

Cardiomyocytes were differentiated using a monolayer method as previously described (Feyen et al. 2021). The iPSCs were seeded in 6-wells at a density of 1.2 × 10^5^ cells per well and grown for four days prior to differentiation. Differentiation was initiated by replacing the E8 media with RPMI supplemented with B27 without insulin (A1895601, Life Technologies) and 6 µM CHIR-99021 (CT99021, Selleckchem). Two days later the media was replaced with RPMI supplemented with B27 without insulin. Cultures were then treated with 3 µM IWR-1 (I0161, Sigma) in RPMI supplemented with B27 without insulin for two days. The cultures were then maintained in RPMI with B27 with insulin (17504-044, Life Technologies) and glucose starved for three days (using RPMI minus glucose). After glucose starvation, iPSC-CMs were maintained in RPMI with B27. Cells were collected at specific time points during differentiation, day 0 (iPSC), day 2 (i-Mes), day 5 (i-CP), day 15 (i-pCM), and day 30 (i-CM), The cells from three independent differentiation batches for each time point were collected and pooled for scATAC analysis.

### Endothelial cells differentiation

The iPSCs were cultured as described above until reaching 80% confluence. The medium was switched to RPMI-B27 without insulin (Life Technologies) with 6 μM CHIR99021 for 2 days and then changed to 2 μM CHIR99021 for another 2 days. During differentiation, from days 4 to 12, the medium was changed to EGM2 (Lonza) supplemented with vascular endothelial growth factor (VEGF) (50 ng/ml) (PeproTech), bone morphogenetic protein 4 (BMP4) (20 ng/ml), and fibroblast growth factor 2 (FGF2) (20 ng/ml) (PeproTech). On day 12, cells were dissociated using TrypLE for 5 min and sorted using CD144-conjugated magnetic microbeads (Miltenyi Biotec) according to the manufacturer’s instructions. CD144-positive cells were seeded on 0.2% gelatin-coated plates and maintained in EGM2 medium supplemented with 10 μM transforming growth factor β (TGFβ) inhibitor (SB431542). (Selleckchem). After passage 2, iPSC-ECs were cultured in EGM2. The iPSC-ECs were analyzed at passage 3 post differentiation.

### Epicardium cells differentiation

The iPSC-derived epicardial cells were differentiated in a chemically defined medium (CDM), which is composed of 50% IMDM, 50% Ham’s F-12 Nutrient Mix, 1% chemically defined lipid concentrate, 2 mM Glutamax, 1 mg/ml PVA, 15 μg/ml transferrin, and 450 μM monothioglycerol. When hiPSCs reached ∼80% confluency they were dissociated with 1 ml of Accutase (Sigma) and replated a density of 1.5 x10^4^ cells/cm^2^ in 6-well plates and cultured in iPS-Brew medium (Miltenyi Biotech) supplemented with 10 μM Y27632. The next day (day 1), each well was washed with D-PBS, and epicardial cells differentiation was initiated by adding the mid-primitive streak induction medium (consisting of 10 ng/ml Activin A, 6 μM CHIR99021, 50 ng/ml BMP4, 20 ng/ml FGF2, and 2 μM LY294002 in CDM). On day 2, each well was refreshed with the lateral plate mesoderm induction medium (consisting of 1 μM A83-01, 30 ng/ml BMP4, and 1 μM C59 in CDM). On days 3-4, each well was refreshed with the splanchnic mesoderm induction medium (consisting of 1 μM A83-01, 30 ng/ml BMP4, 1 μM C59, 20 ng/ml FGF2, and 2 μM retinoic acid in CDM). On days 5-8, the media was refreshed with the septum transversum induction medium (consisting of 2 μM retinoic acid and 40 ng/ml BMP4 in CDM). On day 9, cells were dissociated using Accutase and sparsely seeded (10^4^ cells/cm^2^) on gelatin-coated 6-well plates in the proepicardium induction medium (consisting of 100 μg/ml ascorbic acid, 2 μM of retinoic acid, and 0.7 μg/ml insulin in CDM) for 2 days without medium change. Starting at day 11, each well was refreshed every other day with the epicardial cell induction/maintenance medium (consisting of 100 μg/ml ascorbic acid, 10 μM SB431542, and 0.7 μg/ml insulin in CDM). The iPSC-derived epicardial cells can preserve their cell type-specific markers (e.g., TBX18, WT1, and TCF21) for at least 18 passages in the epicardial cell induction/maintenance medium.

### Cardiac fibroblast differentiation

To generate cardiac-specific fibroblasts, hiPSC-derived epicardial cells were dissociated with Accutase and plated at a density of 10^4^ cells/cm^2^ in 6-well plates and cultured in fibroblast growth medium (Lonza) supplemented with 20 ng/ml FGF2 and 10 μM SB431542. The medium was refreshed every other day for 6 days. When the fibroblasts reached ∼90% confluency, they were dissociated and split at a 1:3 ratio in fibroblast growth medium supplemented with 10 μM SB431542 for long-term maintenance. The differentiated fibroblasts exhibit a quiescent phenotype with negligible (< 5%) α-SMA expression for at least five passages.

### Smooth muscle cell differentiation

To generate cardiac-specific smooth muscle cells (SMCs), iPSC-derived epicardial cells were dissociated with Accutase and seeded at a density of 3×10^4^ cells/cm^2^ were seeded in the nascent SMC induction medium (consisting of 100 μg/ml ascorbic acid, 0.7 μg/ml insulin, 10 ng/ml Activin A, and 10 ng/ml PDGF-BB in CDM) for 2 days. The medium was refreshed every other day with Medium 231 supplemented with SMGS (ThermoFisher) for at least 14 days to allow the expression of SMC-specific markers (e.g., TAGLN, CNN1, SMTNB, and MYH11).

### Single-cell ATAC-seq on iPSC-derived cardiac cells and human fetal heart

The iPSC-derived cardiac cells were dissociated using Tryple Express and resuspended in the RPMI medium. The human fetal hearts were minced and digested using Liberase (Sigma) for 10 min at 37°C, and resuspended in RPMI+B27 medium to stop the enzymatic reaction. The digested tissue was passed through a 70 μm filter before proceeding to single-nuclei sample preparation. Cells with viability > 90% were washed in ice-cold ATAC-seq resuspension buffer (RSB, 10 mM Tris pH 7.4, 10 mM NaCl, 3 mM MgCl2), spun down, and resuspended in 100 µL ATAC-seq lysis buffer (RSB plus 0.1% NP-40 and 0.1% Tween-20). Lysis was allowed to proceed on ice for 5 minutes, then 900 µL RSB was added before spinning down again and resuspending in 50 µL 1X Nuclei Resuspension Buffer (10x Genomics). A sample of the nuclei was stained with Trypan Blue and inspected to confirm complete lysis. Nuclei were processed using a 10X chromium single-cell ATAC-seq kit (V1 version, 10X Genomics) at the Stanford Functional Genomics Facility (SFGF). All samples were sequenced using the Illumina HiSeq 4000 (150 bp paired-end).

### CRISPR–Cas9-mediated genome editing of iPSCs

The genomic region (300-400bp) corresponding to *JARID2* cRE was deleted using CRISPR-Cas9 genome editing. Two guide RNAs (gRNAs) flanking the cRE upstream of *JARID2* were designed using a web-based tool (Benchling) and chosen based on a high score for on-target binding and the lowest off-target score. For cRE deletion, iPSCs (3.5×10^5^) were nucleofected (1200 V, 20 ms, 1 pulse) with 60 pmoles sgRNA (Synthego) and 20 pmoles SpCas9 nuclease (Synthego) using the Neon Transfection System (ThermoFisher Scientific) and the 10 μl tip per the manufacturer’s instructions.). After electroporation, iPSCs were plated in E8 medium supplemented with 5 μM Y-27632 into a 12-well plate coated with Matrigel. After recovering (3 days post electroporation), the cells were dissociated with TrypLE Express and were plated in 6-well plates at a density of 2,000 cells per well. About 10 days after transfection, colonies were picked into 96-well plates and a small proportion of cells from each colony were used for DNA extraction using Quick Extract solution (Epicenter) and direct PCR with Prime STAR® GXL DNA Polymerase (Clontech). PCR amplicons were sequenced by Sanger to verify the deletion (**SFigure 17**).

## Computational methods

### Fetal tissue - scATAC processing

Raw sequencing data were converted to FASTQ format using ‘cellranger-atac mkfastq’ (10x Genomics, v.1.2.0). 150 bp paired-end (PE) scATAC-seq reads were aligned to the GRCh38 (hg38) reference genome and quantified using ‘*cellranger-atac count*’ (10x Genomics, v.1.2.0).

### Fetal tissue - scATAC-seq quality control, dimensionality reduction, and filtering of cell types

Mapped Tn5 insertion sites (fragments.tsv files) from cellranger were read into the ArchR (v0.9.4) R package (Granja et al. 2021). To ensure that each cell was both adequately sequenced and had a high signal-to-background ratio, we filtered cells with fewer than 1,000 unique fragments and enrichment at TSSs below 6. To calculate TSS enrichment, genome-wide Tn5-corrected insertions were aggregated ±2,000 bp relative (TSS-strand-corrected) to each unique TSS. This profile was normalized to the mean accessibility ±1,900–2,000 bp from the TSS, smoothed every 51 bp and the maximum smoothed value was reported as TSS enrichment in R (**SFigure 1**). Latent Semantic Indexing (LSI) dimensionality reduction was computed (*iterations = 4, res = c(0.2,0.2,0.6,0.8), variable features = 25000, dim = 30*) for each specific time point (SFigure 2a,b,c) and repeated the same steps by appending fragment files from all three timepoints together (**SFigure 2d,e**). We did not observe any significant batch effects after the fourth iteration of iterative LSI. We computed chromatin-derived gene accessibility scores by aggregating scATAC-seq reads in each cell weighted by distance from each gene within its cis-regulatory domain (Granja et al. 2021). A preliminary cell-type annotation was performed using these gene accessibility scores of known cell type markers (**Figure 1c,d**, SFigure 3, **Table S3**).

We observed two populations of cell types identified to be macrophages (**SFigure 2f**) and immune cells **(SFigure 2g**). Even though these sets of cell types are of interest from a biological standpoint, they do not directly contribute to the cardiogenesis process and hence were dropped from subsequent analysis. The final UMAP used in all subsequent analyses was generated by repeating the above mentioned iterative LSI with the same parameters as above after removing barcodes corresponding to the macrophage and immune cell clusters (Figure 1c). Final cell-type annotations for each cluster were assigned based on gene accessibility scores of marker genes of known cardiac cell types (**Figure 1c,d**, SFigure 3, **Table S3**).

### Fetal tissue - Peak calling in scATAC-seq datasets

Single-cell chromatin accessibility data were used to generate pseudobulk group coverages based on high-resolution cluster identities of scATAC-seq datasets before peak calling with MACS2 v2.1.1.20160309 (Yong Zhang et al. 2008) using the *addReproduciblePeakSet()* in ArchR (Granja et al. 2021). A background peak set controlling for total accessibility and GC-content was generated using *addBgdPeaks()*. Overlapping peaks were handled using an iterative removal procedure as previously described in (Corces et al. 2018). First, the most significant (MACS2 *q*-value) extended peak summit is kept and any peak that directly overlaps with that significant peak is removed. This process reiterates to the next most significant peak until all peaks have either been kept or removed owing to direct overlap with a more significant peak. The most significant extended peak summits for each cluster were then merged and the previous iterative removal procedure was used. Lastly, we removed any peaks whose nucleotide content had any ‘N’ nucleotides and any peaks mapping to chrY.

### Fetal tissue - scRNA processing

Raw sequencing data from two previous studies (Miao et al. 2020; Cui et al. 2019) corresponding to post-conception week (PCW) 6, 8 and 12, were converted to FASTQ format using the command ‘*cellranger mkfastq*’ (10x Genomics, v.3.1.0). scRNA-seq reads were aligned to the GRCh38 (hg38) reference transcriptome (Ensembl 93) and quantified using ‘*cellranger count*’ (10x Genomics, v.3.1.0). The filtered matrices from cell ranger count were combined with the filtered matrices of other datasets from (Asp et al. 2019) and (Suryawanshi et al. 2020) corresponding to PCW6 and 19 to create the scRNA object.

Count data were further processed using the ‘Seurat’ R package (v.3.1.4) (Stuart et al. 2019), using GENCODE v.27 for gene identification. We removed cells with less than 500 expressed (gene read counts > 0) genes, cells with less than 500 reads, and cells with more than 40% read count corresponding to mitochondrial genes. Genes not contained in the GENCODE annotation were excluded from further analysis. Gene level read count data was scaled to 10,000 (TP/10k) and log_2_ transformed. We performed Principal Component Analysis (PCA) restricting to the 2,000 most variable genes as defined by Seurat. The top 50 principal components (PCs) were used for downstream clustering. Clusters were identified using Leiden clustering implemented in Seurat’s *‘FindClusters()’* function (‘resolution=1’). 2-dimensional representations were generated using uniform manifold approximation and projection (UMAP) (McInnes et al., 2020) as implemented in Seurat and the ‘uwot’ R packages (v.0.1.8; parameter settings: ‘min.dist=0.8’, ‘n.neighbors=50’, ‘cosine’ distance metric). We observed that the clustering was strongly influenced by sample of origin indicating significant batch effects (**SFigure 4a**). To correct these batch effects, we used Harmony (Korsunsky et al. 2019) with max_iters=5 and other parameters set to their default values. We then reran Leiden clustering with the top 30 components from Harmony and generated a 2D UMAP for the corrected data with the same functions listed above. Post harmonization, clusters did not appear to be affected by the sample of origin (**SFigure 4a**). Cell-type annotations for each cluster were assigned based on the expression of known marker genes of cardiac cell types (SFigure 4b,c, **Table S3**).

### Fetal tissue - Matching cells from scRNA-seq and scATAC-seq data

Canonical correlation analysis (CCA) as implemented in Seurat (Stuart et al. 2019) was used to align and match cells from the scRNA-seq and scATAC-seq experiments from each gestational time point individually. For this purpose, we computed log_2_-transformed gene accessibility scores as surrogates for gene expression in the cells profiled by scATAC-seq. As integration features, we used the union of the 2,000 most variable genes in each modality as input to Seurat’s ‘*FindTransferAnchors()*’ function with reduction method ‘cca’ and parameter ‘k.anchor=10’. For each cell profiled by scRNA-seq, we identified the nearest neighbor cell in scATAC-seq by applying nearest-neighbor search in the joint CCA L2 space. Nearest neighbors were determined using the ‘FNN’ R package (https://rdrr.io/cran/FNN/) employing the ‘kd_tree’ algorithm with Euclidean distance. These nearest-neighbor-based cell matches from all gestational time points were concatenated to obtain dataset-wide cell matches across both modalities (**SFigure 4d,e**).

### BPNet deep learning models to predict base-resolution, cell-type resolved pseudo-bulk scATAC-seq profiles from DNA sequence

BPNet is a sequence-to-profile convolutional neural network that uses one-hot-encoded DNA sequence (A=[1,0,0,0], C=[0,1,0,0], G=[0,0,1,0], T=[0,0,0,1]) as input to predict single nucleotide-resolution read count profiles from assays of regulatory activity (Avsec, Weilert, et al. 2021; Trevino et al. 2021). The models take in a sequence context of 2,114 bp around the summit of each ATAC-seq peak and predict cluster-specific scATAC-seq pseudo-bulk Tn5 insertion counts at each base pair for the central 1,000 bp. The BPNet model also uses an input Tn5 bias track which is concatenated to the pre-final layer as explained below. Our BPNet model is a higher capacity version of the architecture introduced in (Avsec, Weilert, et al. 2021). The model architecture consists of 8 dilated residual convolution layers, with 500 filters in each layer. At each layer, the Keras Cropping 1D layer is used to clip out the two edges of the sequence, to match the inputs concatenated to the output of each convolution, which naturally trims the 2,114 bp sequence to a final 1,000 bp profile. Each dilated convolutional layer has a kernel width of 21 and the dilation rate is doubled for every convolutional layer starting at 1. The model predicts the base-resolution 1,000 bp length Tn5 insertion count profile using two complementary outputs: (1) the total Tn5 insertion counts over the 1,000 bp region, and (2) a multinomial probability of Tn5 insertion counts at each position in the 1,000 bp sequence. The predicted (expected) count at a specific position is a multiplication of the predicted total counts and the multinomial probability at that position. To predict the total counts in the 1,000 bp window, the output from the last dilated convolutional layer is passed through a GlobalAveragePooling1D layer in Keras. We estimate the “tn5 bias” for the input sequence using the TOBIAS method (Bentsen et al. 2020). This total bias is concatenated with the output of the pooling layer and passed through a Dense layer with 1 neuron to predict total counts. To predict the per-base logits of the multinomial probability profile output, the output from the last dilated residual convolution is appended with per base TOBIAS “tn5 bias” and passed through a final convolution layer with a single kernel and a kernel width of 1 to predict the per-base logits. BPNet uses a composite loss function consisting of a linear combination of a mean squared error (MSE) loss on the log of the total counts and a multinomial negative log-likelihood loss (MNLL) for the profile probability output. We use a weight of [4.9, 4.3, 18.5, 9.8, 8.9, 4.8, 4.6, 4.9, 12.4, 15.4, 4.3, 6.3, 1.4, 2.6, 7.6, 2.3, 16.3, 7.1 & 3.7] for the MSE loss for clusters c0–c20 (c15-c16 combined as one model), and a weight of 1 for the MNLL loss in the linear combination. The MSE loss weight is derived as the median of total counts across all peak regions for each cluster divided by a factor of 10 (Avsec, Weilert, et al. 2021). We used the ADAM optimizer with early stopping patience of 3 epochs.

A separate BPNet model was trained on pseudobulk scATAC-seq profiles from each scATAC-seq cluster. We used a 5-fold chromosome hold-out cross-validation framework for training, tuning, and test set performance evaluation. The training, evaluation, and test chromosomes used for each fold are as follows. Test chromosomes: fold 0: [chr1], fold 1: [chr19, chr2], fold 2: [chr3, chr20], fold 3: [chr13, chr6, chr22] & fold 4: [chr5, chr16]. Validation chromosomes: fold 0: [chr10, chr8], fold 1: [chr1], fold 2: [chr19, chr2], fold 3: [chr3, chr20] & fold 4: [chr13, chr6, chr22]. The model’s performance was evaluated using two different metrics for the two output tasks separately. For the total counts predicted for the 1,000 bp region, the model’s performance is computed with the Spearman correlation of predicted counts to actual counts. The profile prediction performance is evaluated using the Jensen-Shannon Distance, which computes the divergence between two probability distributions; in this case, the observed and predicted base-resolution probability profile over each 1,000 bp region.

For each cell type, BPNet models were trained, tuned, and evaluated on genomic windows consisting of 1 kb scATAC-seq profiles from (1) signal windows centered at summits of scATAC-seq peaks from the cell type and (2) background windows randomly sampled across the genome such that the number of background windows was 10% of the number of signal windows. The selected signal and background windows were further augmented with upto 10 random jitters (+/- 1000 bp). Code for training BPNet models are available at https://github.com/kundajelab/Cardiogenesis_Repo.

### BPNet model-derived DeepLIFT/DeepSHAP nucleotide contribution scores of accessible cRE sequences

We used the DeepLIFT algorithm (Shrikumar, Greenside, and Kundaje 2017) to interrogate BPNet models and estimate the predictive contribution of each base in any query input sequence to the predicted total counts from the model. DeepLIFT backpropagates a score, analogous to gradients, which is based on comparing the activations of all the neurons in the network for the input sequence to those obtained from neutral ‘reference’ sequences. We use 20 dinucleotide-shuffled versions of each input sequence as reference sequences. We used the DeepSHAP implementation of DeepLIFT (https://github.com/slundberg/shap/blob/0.28.5/shap/explainers/deep/deep_tf.py) to obtain contribution scores for all observed bases in each sequence (Lundberg and Lee 2017b). For each cell type, we obtained consolidated DeepLIFT/DeepSHAP contribution scores for each sequence from each of 5 folds of cross-validation and then averaged the scores per position from the 5 folds.

### Annotation of PWM-based transcription factor motif instances in accessible cREs

We obtained position weight matrix (PWM) models of transcription factor (TF) sequence motifs from the ChromVAR motif catalog called ‘human_pwms_v1’ (Schep et al. 2017), which is collated from the Catalog of Inferred Sequence Binding Preferences (CIS-BP) (Weirauch et al. 2014).

We then annotated PWM-based motif instances in all cRE sequences from all cell types by scanning, scoring, and thresholding (*p*-value < 5e-5) matches from all PWMs using the motifmatchr tool (https://github.com/GreenleafLab/motifmatchr) which uses the MOODSv.1.9.3 library (Korhonen et al. 2009).

### Annotation of cell-type specific active TF motif instances in accessible CREs with high contribution scores and motif mutagenesis scores

For each accessible cRE in each cell type, we defined active motif instances as a subset of PWM-based motif instances that have high DeepLIFT contribution scores or high motif mutagenesis scores from the corresponding cell-type specific BPNet models relative to a null background distribution of corresponding scores.

#### Motif instance contribution scores

We computed the contribution score of each PWM motif instance to accessibility in a specific cell type as the average of the consolidated DeepLIFT contribution scores from the cell-type specific BPNet models over all bases overlapping the motif instance.

#### Motif instance mutagenesis scores

We also inferred mutagenesis scores (motif-ISM) for each PWM-motif instance in a cRE sequence with respect to accessibility in each cell type. To generate the motif-ISM scores for a PWM motif instance in a specific cell type,

1. We first used the fold-0 BPNet model of the specific cell type to predict the total scATAC-seq counts over a 1000 bp window (using a 2114 bp input sequence) centered at the motif instance.
2. We then generated 3 shuffled versions of the input sequence containing the motif instance such that we maintain di-nucleotide frequencies (dinucleotide shuffling).
3. We obtained 3 subsequences overlapping the positions of the original motif instance from the 3 shuffled dinucleotide shuffled sequences.
4. We replaced the subsequence of the motif instance in the original reference sequence with each of the 3 shuffled subsequences.
5. We then use the fold-0 BPNet model to once again predict the total scATAC-seq counts for each of these 3 disrupted sequences containing the shuffled versions of the motif instance.
6. We then computed the log_2_ ratio of the total predicted counts between the reference sequence from step 1. and each of the 3 disrupted sequences from step 5.
7. The motif-ISM score of the instance was computed as the average of the log_2_ ratio score from step 6. over all 3 disrupted sequences.

#### Empirical null distributions

We generated empirical null distributions of motif-instance contribution scores as follows.

1. We constructed dinucleotide frequency preserving shuffled versions of all cREs from from chr4 and chr7.
2. We used the cell-type specific BPNet models from each of the 5 folds to compute DeepLIFT contribution scores over all randomized sequences from step 1. For each sequence, the contribution scores at each base were averaged over all 5 folds.
3. The contribution scores from all bases in all sequences from step 2. were used to derive an empirical null distribution of contribution scores.

We generated empirical null distributions of motif-instance ISM scores as follows.

1. We reused the predicted total scATAC-seq counts for each of these 3 disrupted sequences containing the shuffled versions of the motif instance from step 5. of the motif-ISM estimation process above. We computed the log_2_ ratio of the total predicted counts between each of the 3 pairs of disrupted sequences.
2. The empirical null distribution for motif-ISM scores was derived from the above computed scores over all motif instances in all cRE sequences in chr4 and chr7.

#### Active motif instances

Finally, to identify active motif instances in each cell type, we select PWM-based motif instances that have motif-instance contribution scores or motif-ISM scores that are above the 95th percentile or below the 5th percentile of corresponding empirical null distribution scores of that cell type. All other PWM-based instances were labeled as “inactive”.

### Enrichment of active motif instances and all PWM-motif instances in differential, cell-type specific scATAC-seq peaks

We identified differentially accessible, cell-type specific “marker peaks” for the ventricular cardiomyocyte cluster (vCM) relative to all other clusters using the *getMarkerFeatures()* function in ArchR (Granja et al. 2021), which uses the Wilcoxon Ranksum test to identify marker peaks while controlling for the TSS enrichment and log_10_(unique fragments) of cells when sampling the background set of cells.

We then calculated the Fisher’s Exact test implemented in the *peakAnnoEnrichment()* function in ArchR to compute the enrichment of active motif instances of all TFs expressed in vCMs in vCM marker peaks relative to all vCM peaks. We compute analogous enrichments of all PWM-based motif instances. We compare the statistical significance of enrichments of active and all PWM instances in **Figure 2h,i** and **SFigure 5**.

### ChromVAR motif deviation scores

To compute ChromVAR motif deviation scores for any peak set, a background peak set controlling for total accessibility and GC-content was generated using *addBgdPeaks()* for each cluster in ArchR. Chromvar (Schep et al. 2017) was run with *addDeviationsMatrix()* using active TF motif instances in both peak sets to calculate enrichment of chromatin accessibility over all active motif instances of each TF at single-cell resolution. We then computed the GC-bias-corrected deviation scores using the chromVAR ‘deviationScores’ function used in the *addDeviationsMatrix()* function in ArchR.

### Defining cell transitions and trajectories from scATAC-seq data using optimal transport

#### Computing gene signatures

We created a cell by gene score matrix that was used for computing the gene signatures associated with cell cycle and apoptosis for optimal transport analysis. We used the list of curated genes for cell cycle and apoptosis as suggested in the original optimal transport paper (Schiebinger et al. 2019). We scored cells based on their chromatin derived gene accessibility scores (Granja et al. 2021) of genes in the curated gene signatures. We used the same procedure as in the original manuscript. For each cell, we compute the *z*-score of the gene accessibility scores for each gene in the set. We then clip these *z*-scores in the range of -5 to 5. We define the signature score of the cell to be the mean *z*-score over all genes in the gene set (**SFigure 6a and b**). We estimated the initial growth rate with the same calculations as performed in the original method (Schiebinger et al. 2019) with the cell cycle and apoptosis signal computed from the gene score matrix (**SFigure 6c**).

#### Using gene score matrix for Optimal transport calculation

We performed optimal transport-based trajectory analysis by following the original codebase (https://broadinstitute.github.io/wot/tutorial/) (Schiebinger et al. 2019). The two changes between the original method and our implementation are the use of gene accessibility scores to compute the gene signatures and the use of the cell by gene-accessibility score matrix for inferring the optimal transport maps as compared to the cell by gene expression used in the original method. The cell by gene accessibility score matrix was scaled to read per 10K and log_2_-transformed. The top 2000 variable genes based on Seurat (*FindVariableGenes()* method=”vst”) were retained for further analysis. The coupling inference was obtained using parameters *e = 0.05; l1 = 1; l2 = 50; growth_iters = 3 (Schiebinger et al. 2019)*. We first computed the transport matrices between successive timepoints, inferred long-range temporal couplings and then computed the fate matrices to obtain the transition table (Figure 3b).

#### Chromatin and gene expression dynamics across trajectories

For all the dominant trajectories identified using optimal transport, we identified the clusters that are predicted to be in the trajectory using the transition table (**Figure 3e,f,l,m, SFigure 7,8**). We provided these sets of cells to ArchR’s (Granja et al. 2021) *addTrajectory()* function and assigned cells pseudotime values. We then used a modified version of the *plotTrajectory()* function to plot the chromatin peak dynamics associated with the identified trajectory. We estimated correlation between TF gene expression from scRNA-seq projected into the scATAC-seq subspace and TF ChromVAR deviation scores using *correlateMatrices()* in ArchR (Granja et al. 2021). We defined correlated TFs for each trajectory as those who had correlation values > 0.5.

### iPSC derived *in vitro* cardiac cell types - scATAC-seq data processing, quality control, dimensionality reduction and motif annotations

Raw sequencing data were converted to FASTQ format using ‘*cellranger-atac mkfastq*’ (10x Genomics, v.1.2.0). 150 bp paired-end (PE) scATAC-seq reads were aligned to the GRCh38 (hg38) reference genome and quantified using ‘*cellranger-atac count*’ (10x Genomics, v.1.2.0). Barcode filtering, dimensionality reduction and identification of cell types, and peak calling were done similar to the fetal tissue scATAC-seq data processing (**SFigure 9, 10, 11 & 12**).

PWM-based motif instances for CISBP motifs (Weirauch et al. 2014) were identified in all cell-type resolved scATAC-seq peak regions using *addMotifAnnotations()* in ArchR(Granja et al. 2021) and chromVar (Schep et al. 2017) deviations were computed on this matrix using the *addDeviationsMatrix()* in ArchR (Granja et al. 2021). Differential peaks (**SFigure 12a**) and motif enrichment on the differentially accessible peaks (**SFigure 12b**) for *in vitro* data were obtained using similar methods outlined for the fetal tissue data.

### iPSC derived *in vitro* cardiac cell types - scRNA-seq processing, quality control, dimensionality reduction

Filtered cell x gene matrices of scRNA-seq counts were obtained from (Friedman et al. 2018). The counts matrix was processed in a manner identical to fetal scRNA processing. All functions listed above in Seurat were used again for dimensionality reduction, clustering, and cell-type identification. We observed batch effects with the published scRNA data with cells clustering based on their timepoints instead of their cell types identified in the original manuscript (Friedman et al. 2018). To correct this batch effect, we ran Harmony to correct for time point of origin and obtained the corrected scRNA-seq data and corrected clusters and cell type annotations (**SFigure 13**).

### iPSC derived *in vitro* cardiac cell types - Matching cells from scRNA-seq and scATAC-seq data

CCA based matching of scATAC and scRNA data from the *in vitro* differentiation was done in the same manner as the fetal tissues and the nearest neighbor’s gene expression profile in scRNA was imputed to the scATAC ArchR object (Granja et al. 2021).

### Projecting iPSC derived *in vitro* cardiac cells based on scATAC-seq into the the fetal heart scATAC-seq manifold

We projected the iPSC derived *in vitro* cardiac cells based on the scATAC-seq profiles into the scATAC-seq LSI subspace of fetal heart cells following the procedure described previously (Granja et al. 2019). Briefly, when computing the TF-IDF transformation on the fetal samples, we stored the colSums, rowSums, and SVD. To project cells from additional samples into this subspace, we first zeroed out rows based on the initial TF-IDF rowSums. We next calculated the term frequency by dividing by the column sums and computed the inverse document frequency from the previous TF-IDF transformation. These were then used to compute the new TF-IDF. The resulting TF-IDF matrix was projected into the previously defined SVD of the fetal heart LSI.

### Identifying scATAC-seq peaks across all *in vivo* and *in vitro* cardiac cells

To enable the comparison of epigenomic features between the *in vivo* and *in vitro* cells, we built a combined ArchR object of all post filtered cells from the three fetal heart samples and all the samples from the iPSC differentiation to major cardiac cell types. We performed peak calling on the combined data using ArchR frameworks, as described above. We used these peak calls from the combined object for all the downstream differential analyses between the *in vivo* and *in vitro* nearest cells identified by the projection analysis.

### Identifying differential scATAC-seq peaks and TF motif enrichments between matched *in vivo* and *in vitro* cardiac cell types

To identify differential scATAC-seq peaks between each of the *in vivo* fetal heart cell types and their matched *in vitro* counterparts, we create a combined ArchR object of both the 3 fetal samples and all timepoints from the *in vitro* differentiation study. Peak calling was performed on this combined object as described above and PWM-based motif instances (Weirauch et al. 2014) were used to compute TF motif enrichments and ChromVar deviations as described above.

Differential peaks between *in vivo* and *in vitro* cell types were identified within this integrated peak set. For each pair of match cell types, we obtained the integrated cell x peak matrix. We then computed row-wise two-sided *t*-tests for each peak and estimated the FDR using *p.adjust(method = “fdr”)*. Peaks with absolute log_2_(fold changes) > 0.5 and FDR < 0.05 were labeled as differential.

We next identified the TF motifs enriched in up or down regulated differential peaks relative to all peaks for each pairwise comparison using a Fisher’s Exact test.

### Identifying peak-to-gene links using *in vivo* and *in vitro* data

We computed peak-to-gene links based on correlation between peak scATAC-seq and gene scRNA-seq signals separately for *in vivo* and in vitro cells. We used all cells from *in vivo* samples. For *in vitro* samples, we restricted to cells from terminal cell types which had well-defined *in vivo* counterparts. First, we added the integrated scATAC-seq peaks across *in vitro* and *in vivo* cells to each of these objects using the *addPeakSet()* function in ArchR. Next, we restricted scRNA-seq analysis to a subset of genes that are present in the filtered gene lists of both the *in vivo* and *in vitro* the scRNA-seq objects. We then used CCA as described above to match cells between the scRNA-seq and scATAC-seq experiments for the *in vivo* and *in vitro* samples separately. We identified correlation based peak-to-gene links using the *addPeak2GeneLinks()* function in ArchR (Granja et al. 2021) for the *in vivo* and *in vitro* cells separately. We used a correlation threshold of 0.45 to define linked peak-gene pairs. We compared peak-to-gene links from the *in vivo* and *in vitro* samples to identify peak-gene pairs that were shared and those that were exclusive to each set (**SFigure 14**).

### Enrichment of GWAS traits using stratified linkage disequilibrium score regression (S-LDSC)

We assessed the enrichment of heritability for several cardiovascular diseases and traits in accessible elements stratified linkage disequilibrium score regression (S-LDSC) (Finucane et al. 2015).. We ran LD score regression using the baselineLD_v1.1 model using the 1000G_EUR_Phase3_baseline file (downloaded from https://data.broadinstitute.org/alkesgroup/LDSCORE/; on each cell-type specific peaks, defined as variants in 1000 Genomes with minor allele count >5 in 379 European samples), comparing to summary statistics for atrial fibrillation (Roselli et al. 2018), coronary artery disease (van der Harst and Verweij 2018), heart failure (Shah et al. 2020), and inflammatory bowel disease (Liu et al. 2015). Summary statistics were located using the Cardiovascular Disease Knowledge Portal (https://cvd.hugeamp.org/, accessed August 2020)

### Predicting mutation impact scores of *de novo* non coding mutations from CHD cases and controls on cell-type resolved scATAC-seq profiles using neural network models

We obtained *de novo,* non coding mutations from CHD patients from the Pediatric Cardiac Genomics Consortium (PCGC) and from healthy controls (unaffected siblings) from the Simons simplex collection (SSC) from (Richter et al. 2020). We restricted all analysis to single-nucleotide (point) mutations within these cohorts.

For each cell type, we used cell-type specific BPNet models to predict the allelic impact of all mutations that were found within 1000 bp windows around summits of scATAC-seq peaks in that cell type. For each mutation, we used the BPNet model to predict the base-resolution read count profile corresponding to the input sequence (2,114 bp) containing the reference allele of the mutation at its center. We then used the model to predict the 1 kb base-resolution read count profile (which is decomposed into total predicted counts over 1 kb and base-resolution read probability profile) corresponding to the input sequence (2,114 bp) containing the alternate allele of the mutation at its center. Using these predicted read probability profiles from the two alleles, we computed the impact score of the mutation as the log_2_ fold change in cumulative probability between the reference allele and the alternate allele, over a 100 bp window around the mutation using the formula:

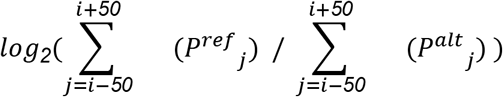

where

*i*= position of mutation

*P^ref^_j_* = predicted profile probability at position *j* for sequence containing reference allele

*P^alt^_j_* = predicted profile probability at position *j* for sequence containing alternate allele

For each mutation, the cell-type specific impact scores were computed and averaged over cluster-specific BPNet models trained on each of 5 folds.

We also computed an alternate mutation impact score based on the predicted cumulative read counts over the 100 bp window around the mutation, instead of the predicted cumulative read probability.

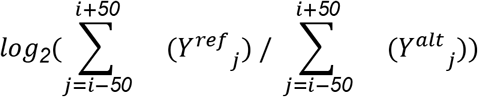

where

*i*= position of mutation

*Y^ref^_j_* = predicted counts at position *j* for sequence containing reference allele

*Y^ref^_j_* = predicted counts at position *j* for sequence containing alternate allele

We found high concordance of cell type specific enrichments of high impact mutations in cases vs. controls for both scores (**SFigure 15c**).

### Thresholding mutation impact scores to define high impact prioritized mutations

Because we are investigating a cohort of children with CHD born to parents without CHD, our expectation is that the majority of these cases will be caused by *de novo* mutations. On average, each individual has approximately 70 such mutations (Richter et al. 2020), and because we assume mutations that lead to CHD are generally rare, we would expect just one would be a causal presentation and we would expect only a fraction of the cohort to have such causal mutations. Based on the expectation that a small proportion of mutations from CHD cases in cell type resolved scATAC-seq peaks will have a causal role, we prioritized **high-impact mutations** in each cell type, as those that have an impact score > 95^th^ percentile of the distribution of cell-type specific impact scores of all mutations from the CHD cohort that fall in 1kb scATAC-seq peak regions in that cell type. The same thresholds were used for mutation impact scores of control mutations as well to obtain enrichments as specified below.

### Selection of prioritized mutations in arteries for deeper investigation

We further restricted deeper investigation into a subset of higher confidence CHD mutations prioritized by the arterial endothelial cells (aEC) BPNet model to those that were within 200 bp (+/- 100 bp) of summits of aEC scATAC-seq peaks that had > 75 reads in a +/- 250 bp window around mutation. For each of these selected mutations, we obtained predicted profiles for sequences centered at the mutation for both alleles as well as the corresponding DeepLIFT scores and active motif instances. The gene closest to the mutation in linear genomic sequence was assigned as the putative target gene of the mutation.

### Cell-type specific enrichment analysis of prioritized mutations in cases relative to controls

To compute the enrichment of case vs. control mutations in scATAC-seq peaks (cREs) of each cell type in the fetal heart, we computed a 2 x 2 contingency table. The first axis splits all *de novo* mutations based on whether they were found in cases versus controls. The second axis splits all *de novo* mutations based on whether they overlap a cluster-specific peak. The enrichment *p*-value and odds-ratio (OR) was computed using the Fisher’s Exact Test implemented in the SciPy package in Python.

We used a similar procedure to estimate enrichment of *de novo* mutations prioritized by cell-type specific models from cases versus control. In this case, the first axis of the 2 x 2 contingency table splits all *de novo* mutations based on whether they were found in cases versus controls. The second axis splits all *de novo* mutations based on whether they are predicted to have a high impact score (> 95^th^ percentile) or not using a cell-type specific BPNet model. High impact score mutations are pre-filtered to those in peak regions in the cell type. This analysis was performed for each cell type separately and for the pseudobulk of all cell types.

### Enrichments of case and control mutations using mutation impact scores from the HeartENN model

We obtained mutation impact scores as computed by the authors of the HeartENN model for all non-coding *de novo* mutations in the PCGC case and SSC unaffected controls (Richter et al. 2020). We retained the *de novo* mutations that overlap 1 kb scATAC-seq peak regions in any of the fetal heart cell types. Finally, we performed Fisher’s exact test for enrichment of high impact (scores >= 0.1 as recommended in (Richter et al. 2020)) mutations in peaks in cases vs controls.

## Supplementary figure legends

**SFigure 1:**
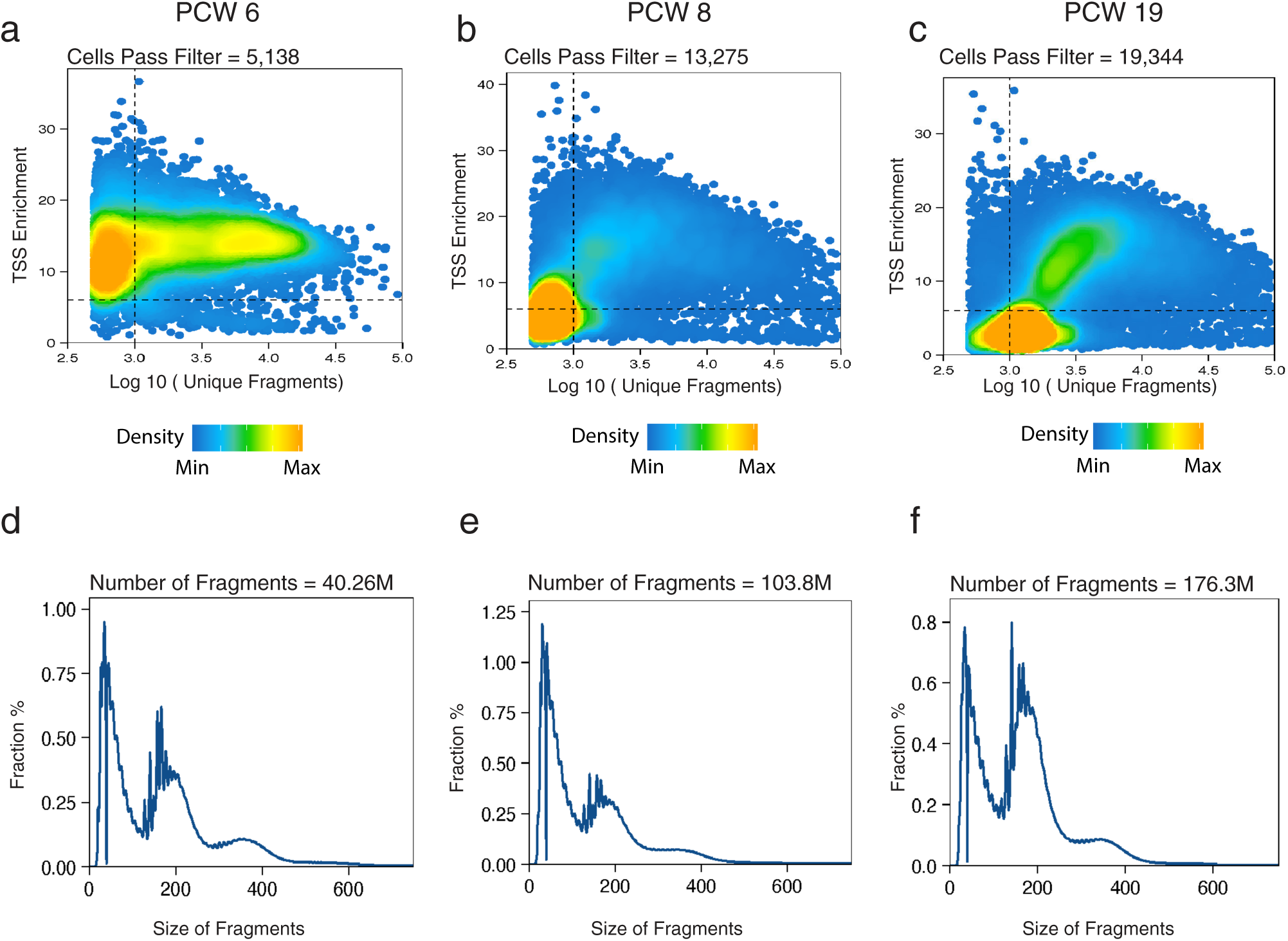
Quality control for scATAC-seq data from fetal hearts at PCW 6 (left), PCW 8 (middle) & PCW19 (right). (a,b,c) Shown are the number of unique ATAC-seq nuclear fragments in each single cell (each dot) compared to TSS enrichment of all fragments in that cell. Dashed lines represent the thresholds for filtering cells (1,000 unique nuclear fragments and TSS score >= 6). (**d, e & f**) The fragment length distribution for PCW 6 (left), PCW 8 (middle) & PCW19 (right).

**SFigure 2:**
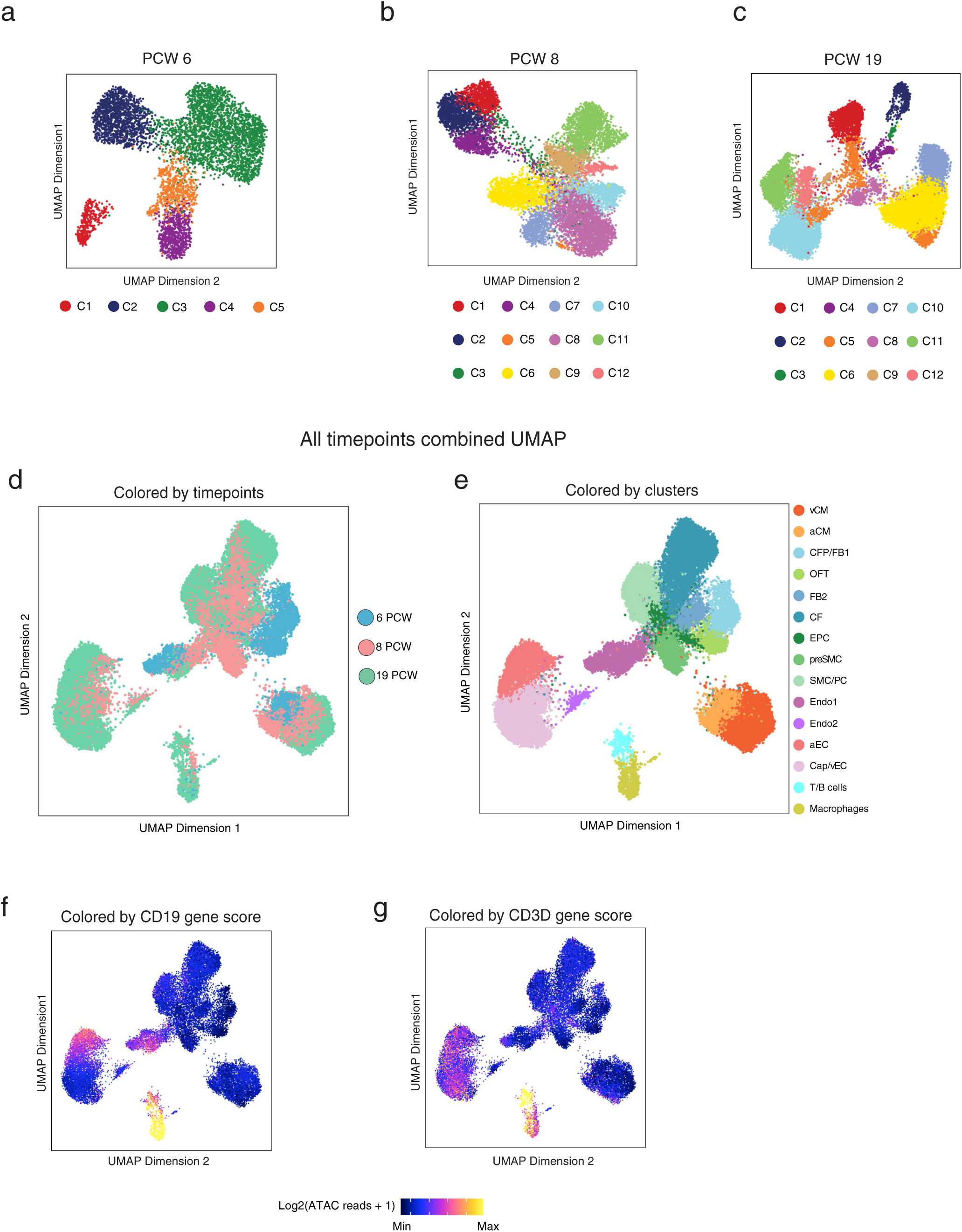
Clustering of cells from scATAC-seq data. (**a, b, c**) Uniform Manifold Approximation and Projection (UMAP) of cells based on open chromatin regions (scATAC-seq) of PCW 6 (left), PCQ 8 (middle), and PCW19 (right) samples. Cells are colored according to clusters. (**d & e**) UMAP of cells from three timepoints combined. Cells are colored according to (**d**) sample gestational time and (**e**) cluster membership. (**f,g**) scATAC-seq gene activity profiling of immune marker genes (**f**) CD19 and (**g**) CD3D.

**SFigure 3:**
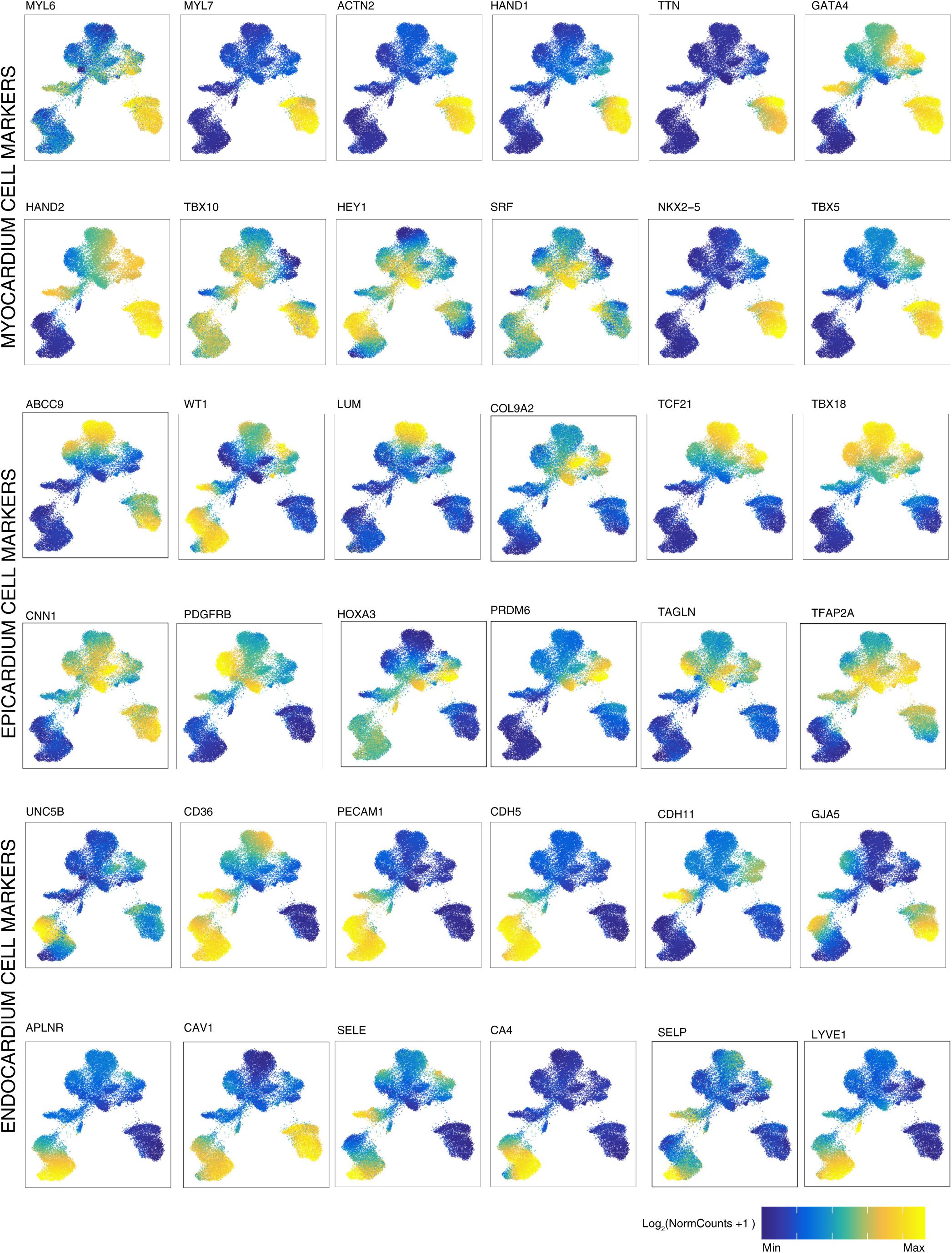
Gene scores of representative cell-type specific marker genes in UMAP of cells based on scATAC-seq data. Units: log_2_(normalized ATAC gene-score). Scale: *MYL6* (min=0.6,max=1), *MYL7* (min=0.25,max=1.4), ACTN2 (min=0.2,max=1.2), *HAND1* (min=0.4,max=1.2),*TTN* (min=0.4,max=2.2),*GATA4* (min=0.5,max=1.6), *HAND2* (min=0.5,max=1.75),*TBX10* (min=0.2,max=0.8),*HEY1* (min=0.8,max=1.4), *SRF* (min=1,max=1.3), *NKX2-5* (min=0.5,max=2),*TBX5* (min=0.2,max=1), *ABCC9* (min=0.15,max=0.7), *WT1* (min=0.4,max=1),*LUM* (min=0.05,max=0.3), *COL9A2* (min=0.2,max=0.6), *TCF21* (min=0.2,max=1),*TBX18* (min=0.2,max=0.9), *CNN1* (min=0.2,max=0.6), *PDGFRB* (min=0.4,max=1.4),*HOXA3* (min=0.2,max=0.8), *PRDM6* (min=0.2,max=1), *TAGLN* (min=0.2,max=0.9),*TFAP2* (min=0.25,max=0.7), *UNC5B* (min=0.9,max=1.4), *CD34* (min=0.4,max=1.2),*PECAM1* (min=0.25,max=1.25), *CDH5* (min=0.3,max=1.5), *CDH11* (min=0.3,max=1.5),*GJA5* (min=0.2,max=1), *APLNR* (min=0.4,max=1.5), *CAV1* (min=0.2,max=0.8),*SELE* (min=0,max=0.45), CA4 (min=0.3,max=1.1), *SELP* (min=0,max=0.25),*LYVE1* (min=0.1,max=0.6)

**SFigure 4:**
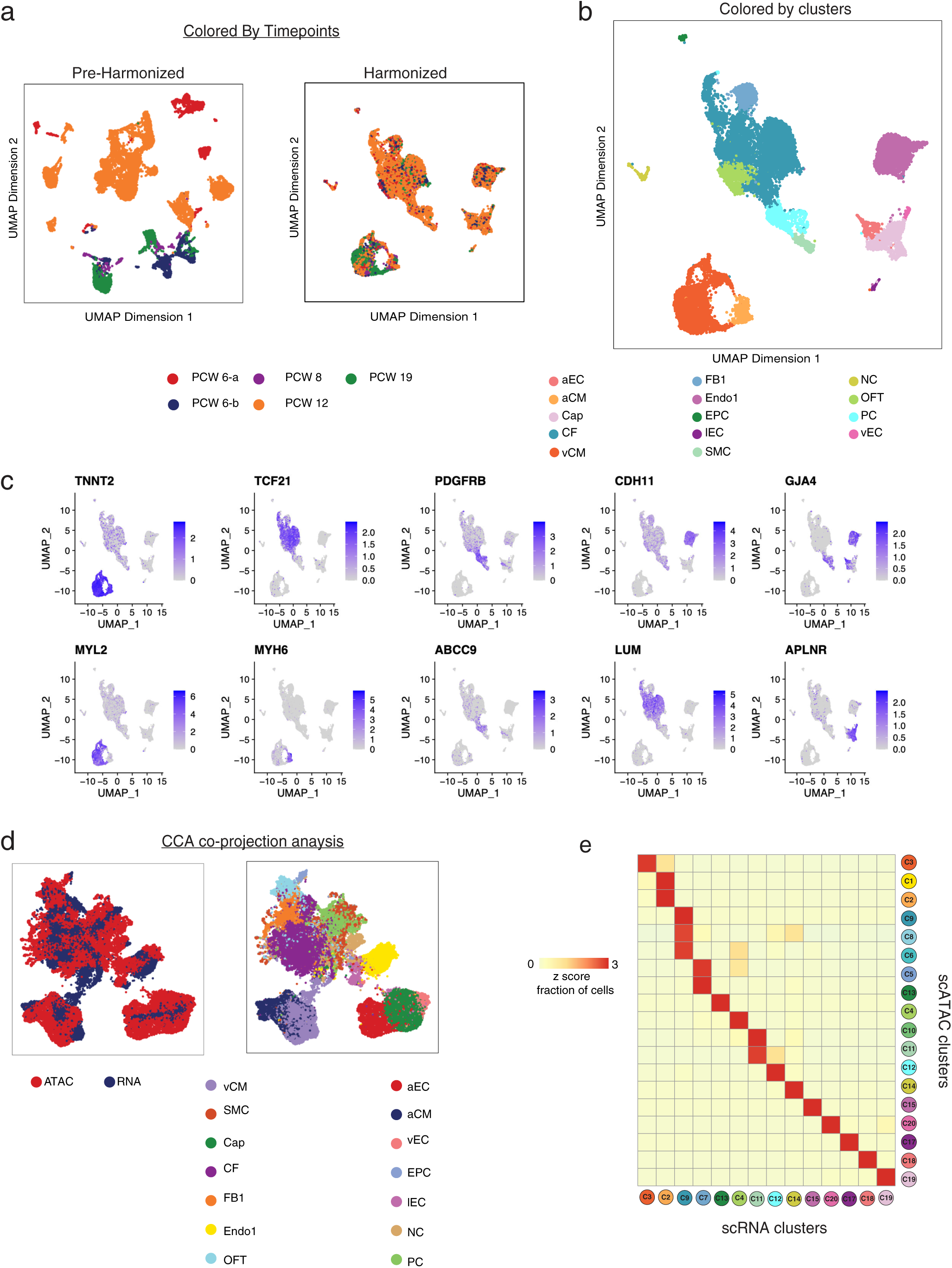
Integration of scRNA-seq & scATAC-seq data using canonical correlation analysis (CCA). **(a)** UMAP of cells from 5 scRNA-seq studies without (left) and with (right) batch effect correction and harmonization using Harmony (right). Cells are colored by the scRNA study of origin. **(b)** Harmonized UMAP of scRNA-seq analysis used for downstream analysis. Cells are colored by clusters. **(c)** Gene expression (Units: TP10K) of cell type specific and cluster specific markers in harmonized scRNA-seq UMAP. **(d)** UMAPs of matched cells from scATAC-seq and scRNA-seq data modalities using the CCA subspace. On the left, cells are colored by their assay type and on the right, cells are colored by clusters from scRNA-seq. **(e)** Heatmap showing the cluster – cluster mapping between scRNA-seq and scATAC-seq clusters after CCA matching.

**SFigure 5:**
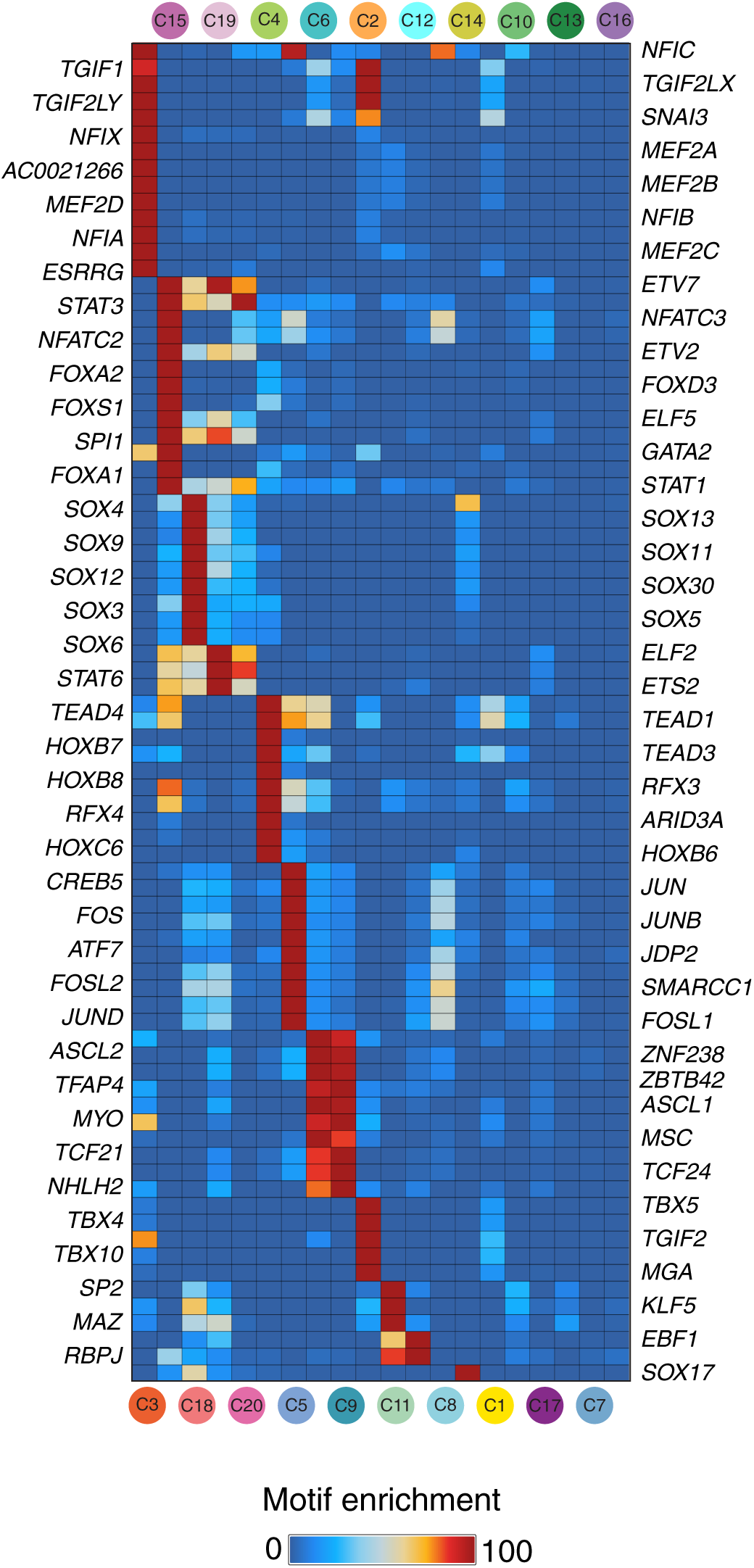
Overlap motif enrichment. Overlap enrichment of position-weight matrix based motif instances in cell-type specific marker scATAC-seq peaks of each cell type cluster from Figure 1e.

**SFigure 6:**
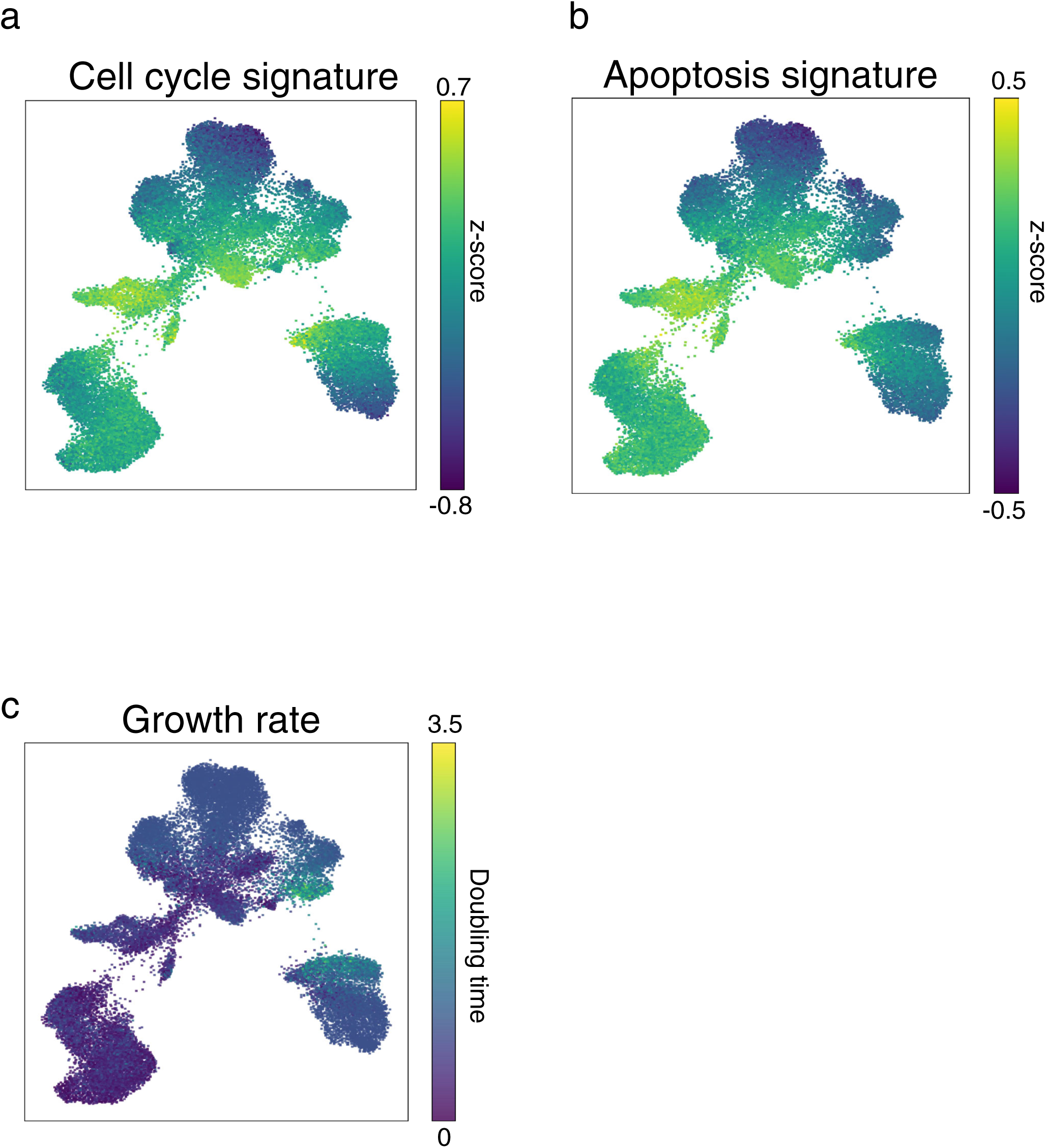
Estimated cell signature scores from optimal transport algorithm applied to gene activity scores from scATAC-seq data. (**a, b, c**) UMAP of cells from scATAC-seq data showing (**a**) cell cycle signature z-scores, (**b**) apoptosis signature z-scores, and (**c**) growth rate estimates

**SFigure 7:**
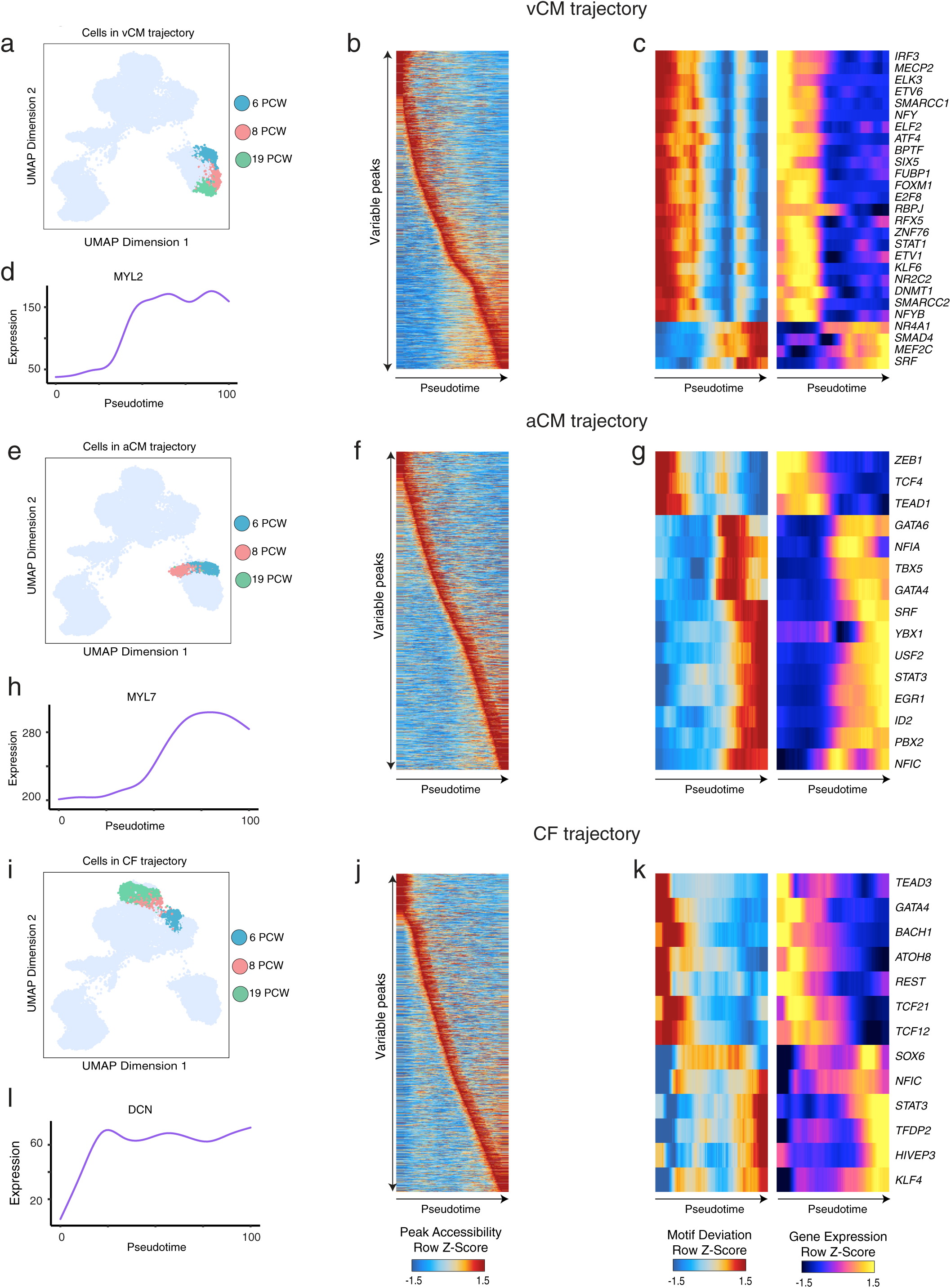
Optimal transport based developmental trajectories for vCM, aCM and CF cells using scATAC-seq. **(a)** UMAPs of scATAC-seq cells in the ventricular cardiomyocyte (vCM) trajectory colored by the gestational sample time. **(b)** Heatmap of scATAC-seq signal (*z*-score of log_2_(RP10K)) of variable peaks identified in the vCM trajectory. **(c)** Heatmaps showing *z*-score of ChromVAR motif deviation scores (left) and gene expression in units of log_2_(TP10K) (right) of TFs with correlated variable activity in cells identified to be in the vCM trajectory, as ordered by pseudotime. **(d)** Expression dynamics of *MYL2*, an important marker gene for the vCM cell type. (**e, f, g, h**) Trajectory analysis for atrial cardiomyocyte cluster (aCM), analysis as above. (**i, j, k, l**) Trajectory analysis for cardiac fibroblast cluster (CF), as above.

**SFigure 8:**
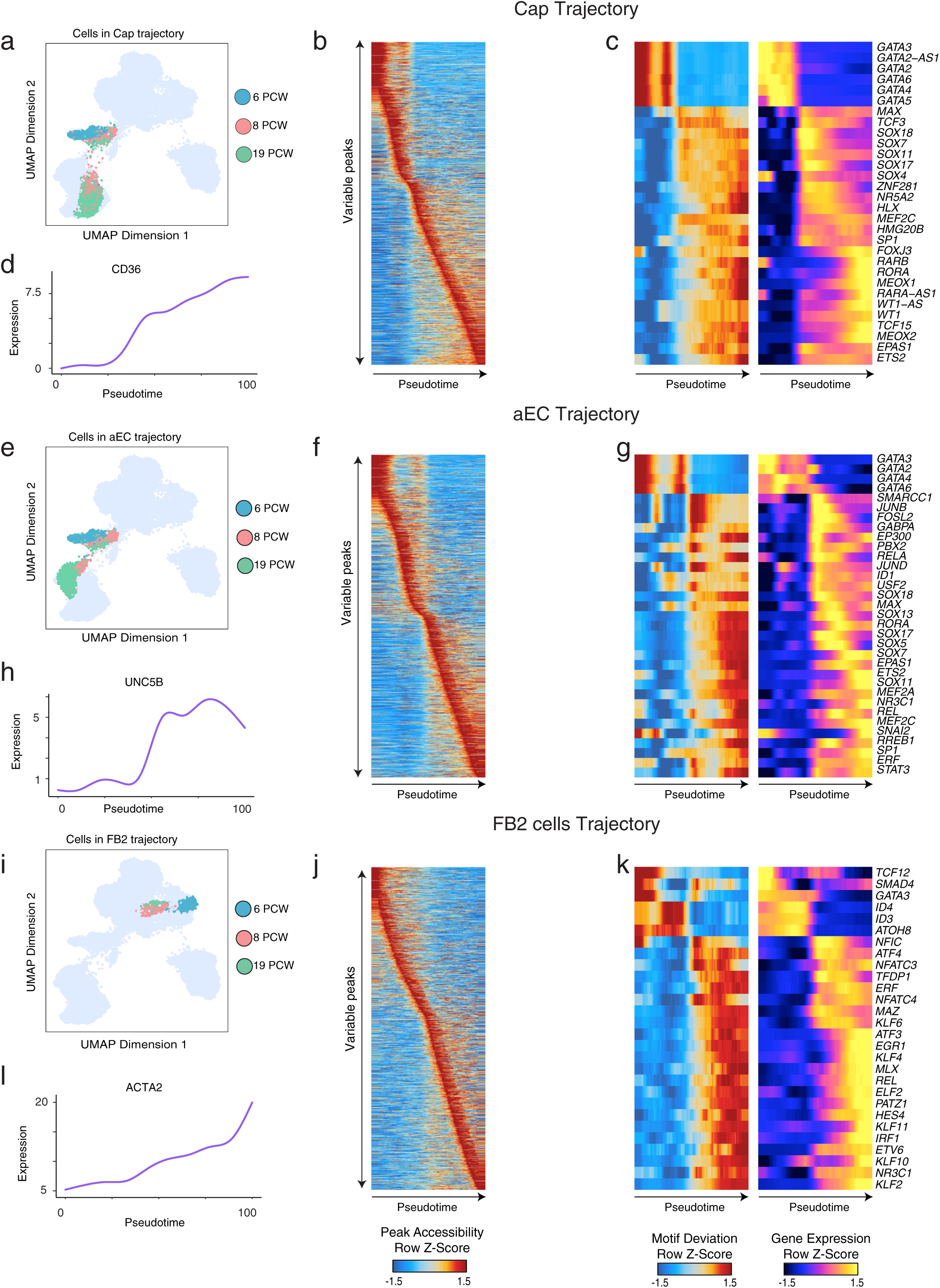
Optimal transport based developmental trajectories for Cap, aEC and FB2 cells using scATAC-seq. **(a)** UMAPs of scATAC-seq cells in the Capillary cells (Cap) trajectory colored by the gestational sample time. **(b)** Heatmap of scATAC-seq signal (*z*-score of log_2_(RP10K)) of variable peaks identified in the vCM trajectory. **(c)** Heatmaps showing *z*-score of ChromVAR motif deviation scores (left) and gene expression in units of log_2_(TP10K) (right) of TFs with correlated variable activity in cells identified to be in the vCM trajectory, as ordered by pseudotime. **(d)** Expression dynamics of *CD36*, an important marker gene for the Cap cell type. (**e, f, g, h**) Trajectory analysis for arterial endothelial cell cluster (aEC), analysis as above. (**i, j, k, l**) Trajectory analysis for Fibroblast like cells 2 (FB2), as above.

**SFigure 9:**
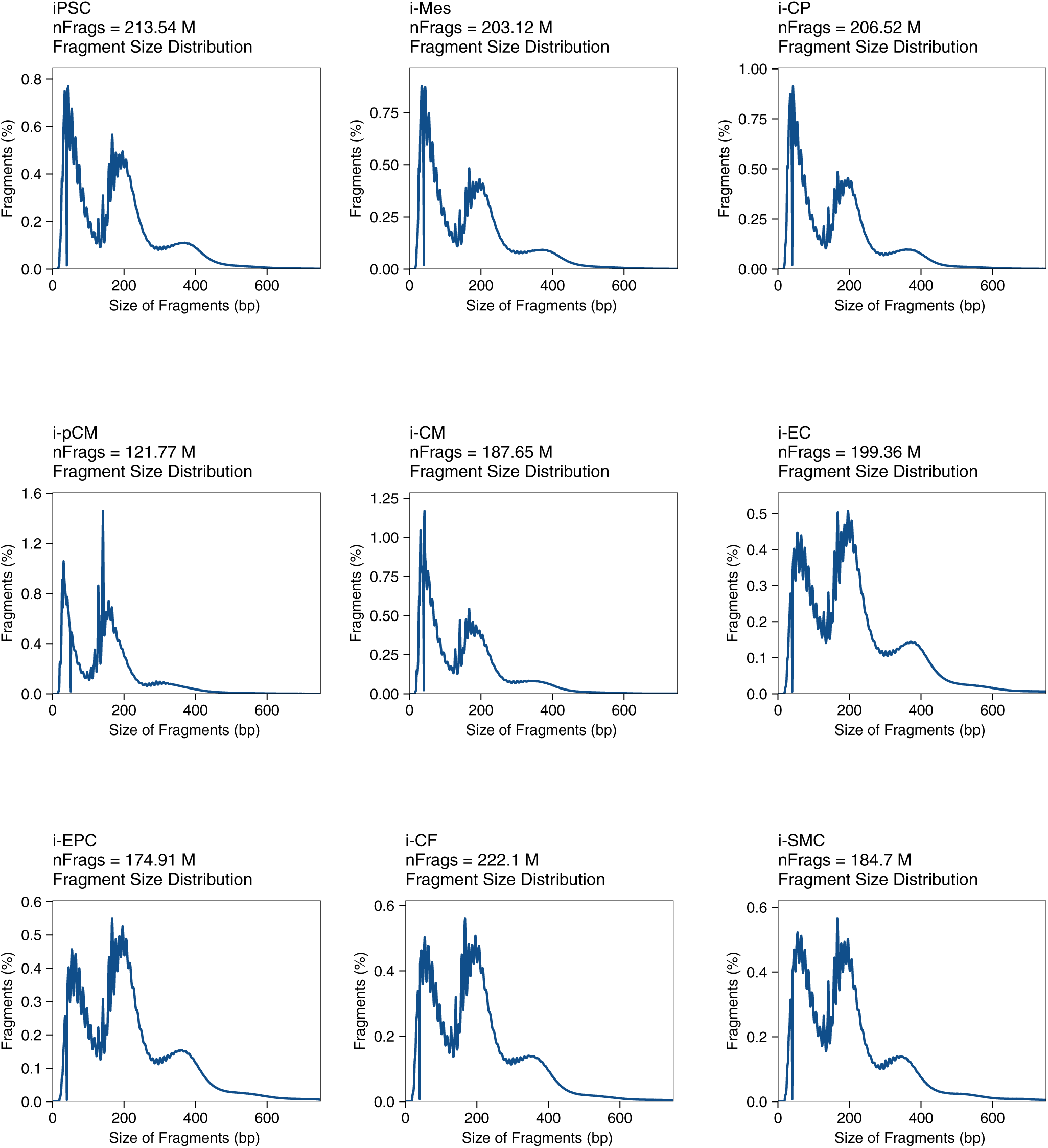
Quality control data for iPS derived cardiac cell types. (**Left to Right, Top to Bottom)** Representative scATAC-seq data quality control filters for Day 0, Day 2, Day 5, Day 15, Day 30 cardiomyocytes, Day 30 endothelial cells, Day 30 epicardial cells, Day 30 cardiac fibroblast cells & Day 30 smooth muscle cells (**top to bottom, left to right**). Shown are the fragment length distribution for the same samples as above.

**SFigure 10:**
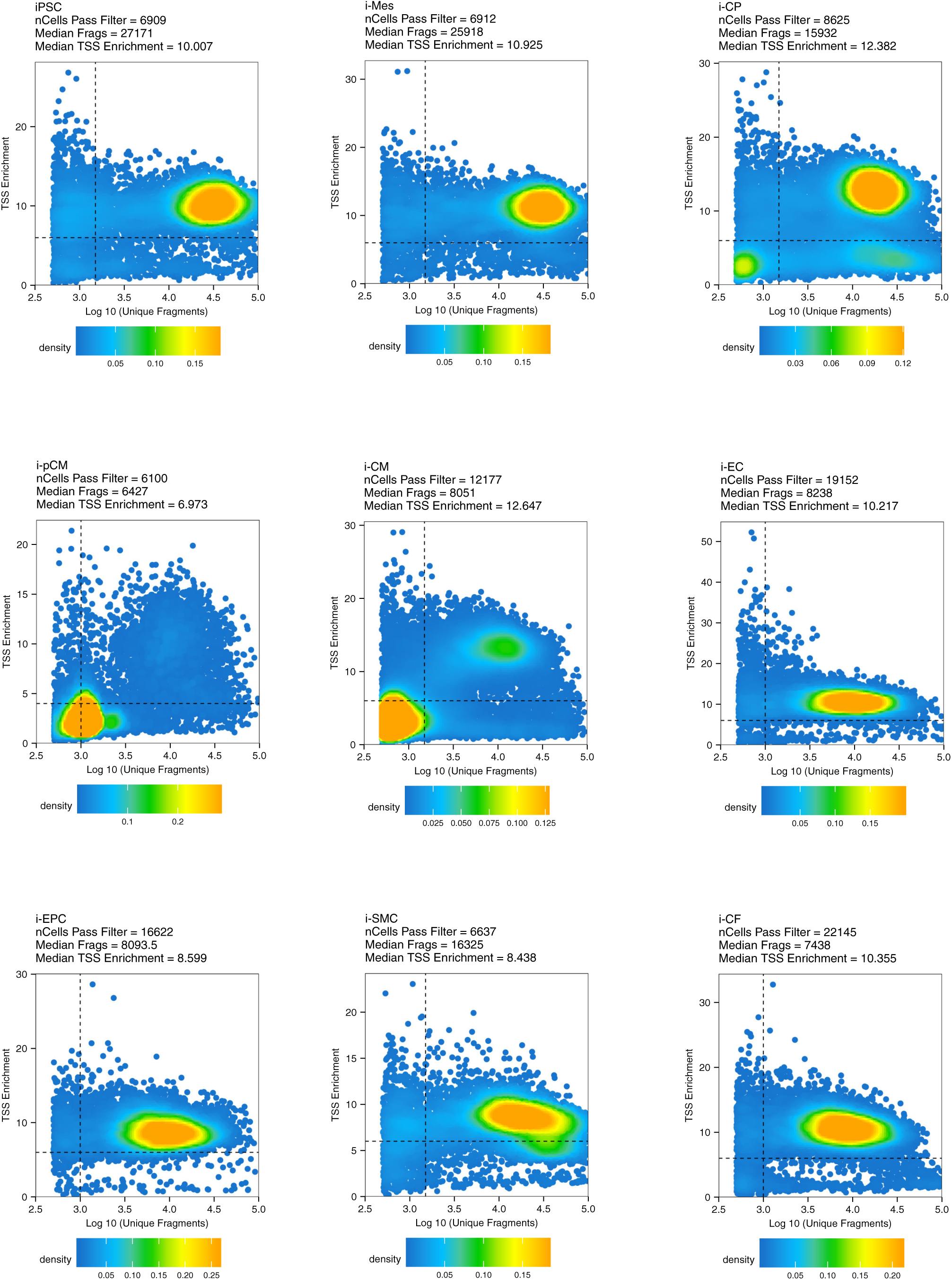
Quality control data for iPS derived cardiac cell types. (**Left to Right, Top to Bottom)** Representative scATAC-seq data quality control filters for Day 0, Day 2, Day 5, Day 15, Day 30 cardiomyocytes, Day 30 endothelial cells, Day 30 epicardial cells, Day 30 cardiac fibroblast cells & Day 30 smooth muscle cells (**top to bottom, left to right**). Shown are the number of unique ATAC-seq nuclear fragments in each single cell (each dot) compared to TSS enrichment of all fragments in that cell. Dashed lines represent the filters for high-quality single-cell data (1,500 unique nuclear fragments and TSS score greater than or equal to 6). (**g, h, i, j, k, l**)

**SFigure 11:**
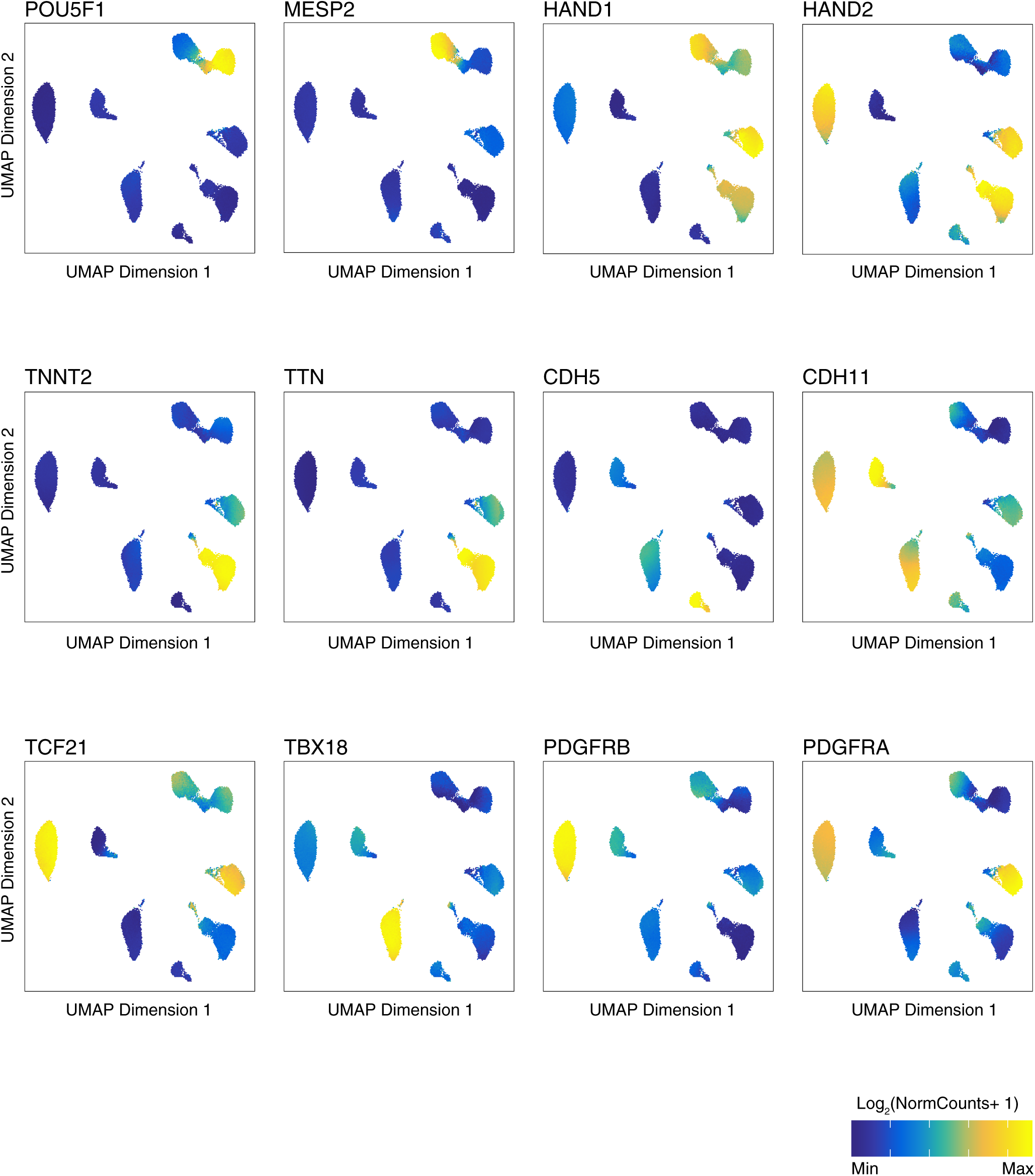
Gene scores of cell type specific markers in iPSC-derived cardiac cell type scATAC-seq data. (**a**) UMAP plots showing gene scores of cell type specific and cluster specific markers.Units: log_2_(normalized ATAC gene-score). Scale: *POU5F1* (min=0,max=0.7), *MESP1* (min=0.25,max=1.25), *HAND1* (min=0.4,max=1.6), *HAND2* (min=0.8,max=1.4), *TNNT2* (min=0.25,max=1.4), *TTN* (min=0,max=2), *CDH5* (min=0.3,max=1.5), *CDH11* (min=0.4,max=1.2), *TCF21* (min=0.2,max=0.9), *TBX18* (min=0.4,max=1), *PDGFRB* (min=0.4,max=1.2) & *PDGFRA* (min=1.4,max=2.2).

**SFigure 12:**
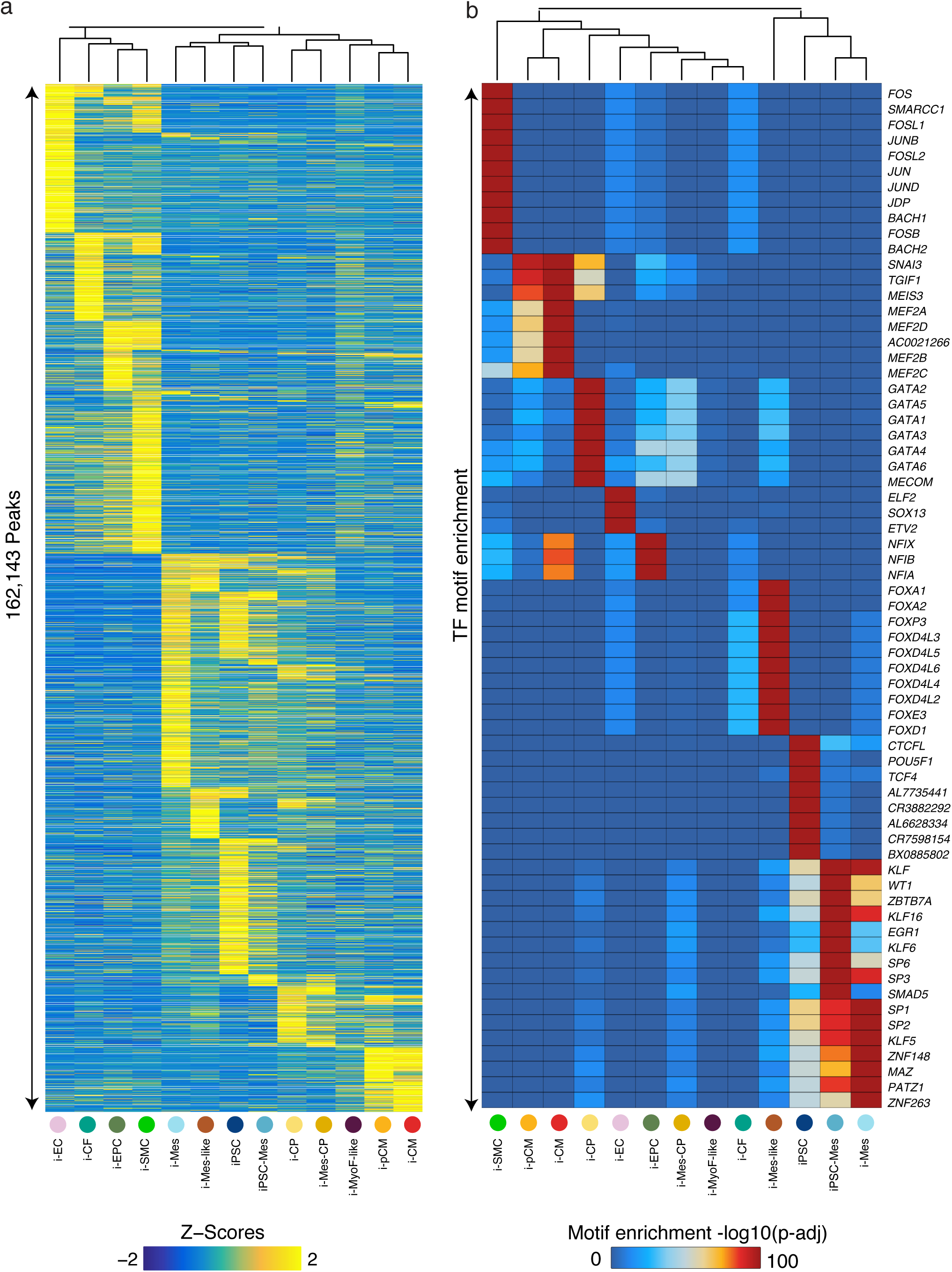
Landscape of variable chromatin accessible peaks and their enriched motifs in iPSC-derived cardiac cell types. **(a)** Heatmap of *z*-scores of log_2_(scATAC-seq read counts) in cREs across iPSC-derived cell type clusters. **(b)** Statistical significance (BH adjusted -log_10_(*p*-value), hypergeometric test) of overlap enrichment of position-weight matrix based motif instances in cell-type specific marker scATAC-seq peaks of each cell type cluster in (**a**).

**SFigure 13:**
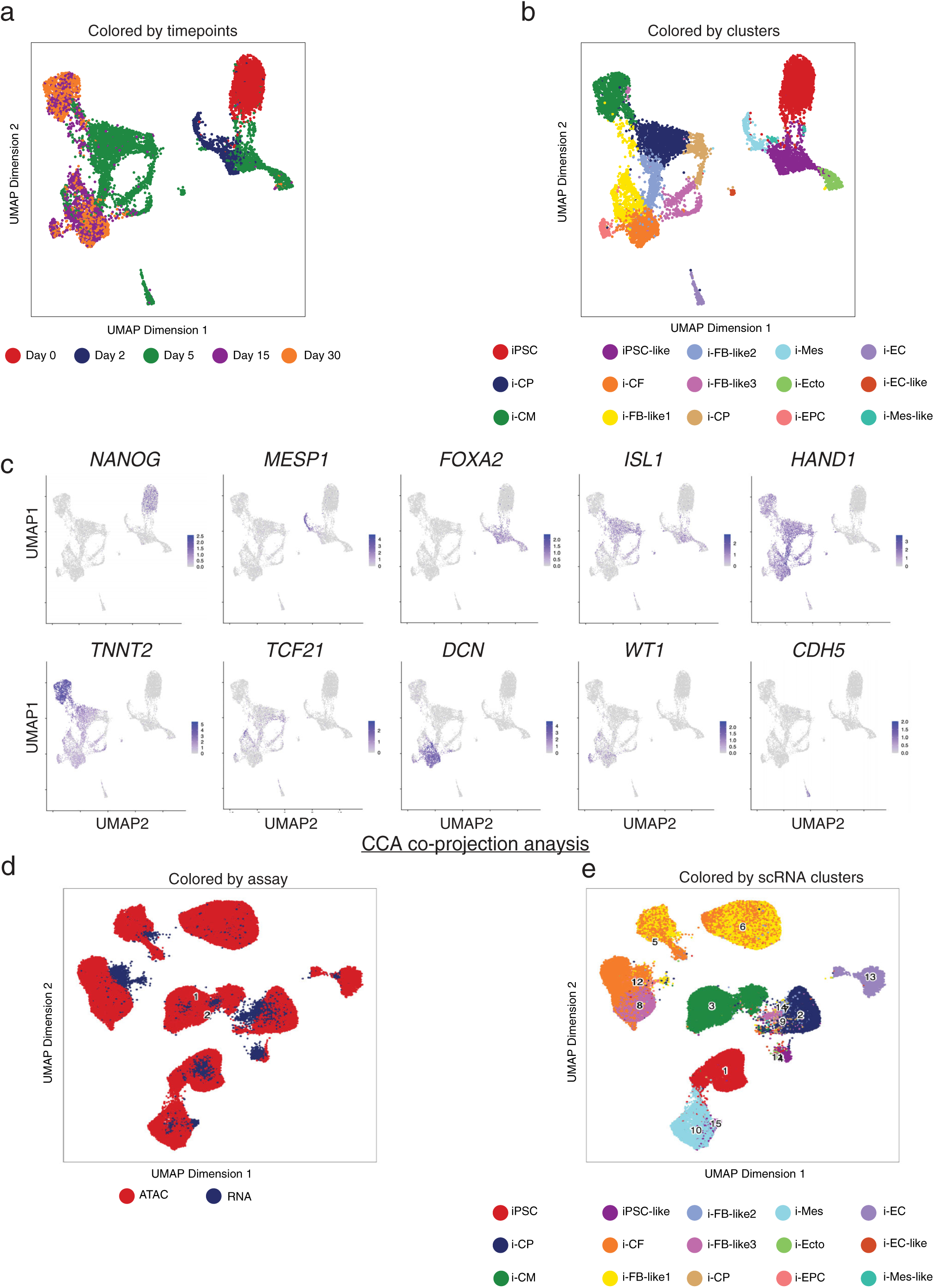
Integration of scRNA-seq & scATAC-seq data from iPSC-derived cell types using canonical correlation analysis (CCA). (**a,b**) UMAP of cells from scRNA-seq in iPSC-derived cardiac cell types. Cells are colored by the differentiation time point (a) and cell-type label of clusters (b). **(c)** Gene expression (Units: log_2_(TP10K)) of cell type specific and cluster specific markers in harmonized scRNA-seq UMAP. (**d,e**) UMAPs of matched cells from scATAC-seq and scRNA-seq data modalities using the CCA subspace, colored by assay type (d) and cell type label of clusters (e).

**SFigure 14:**
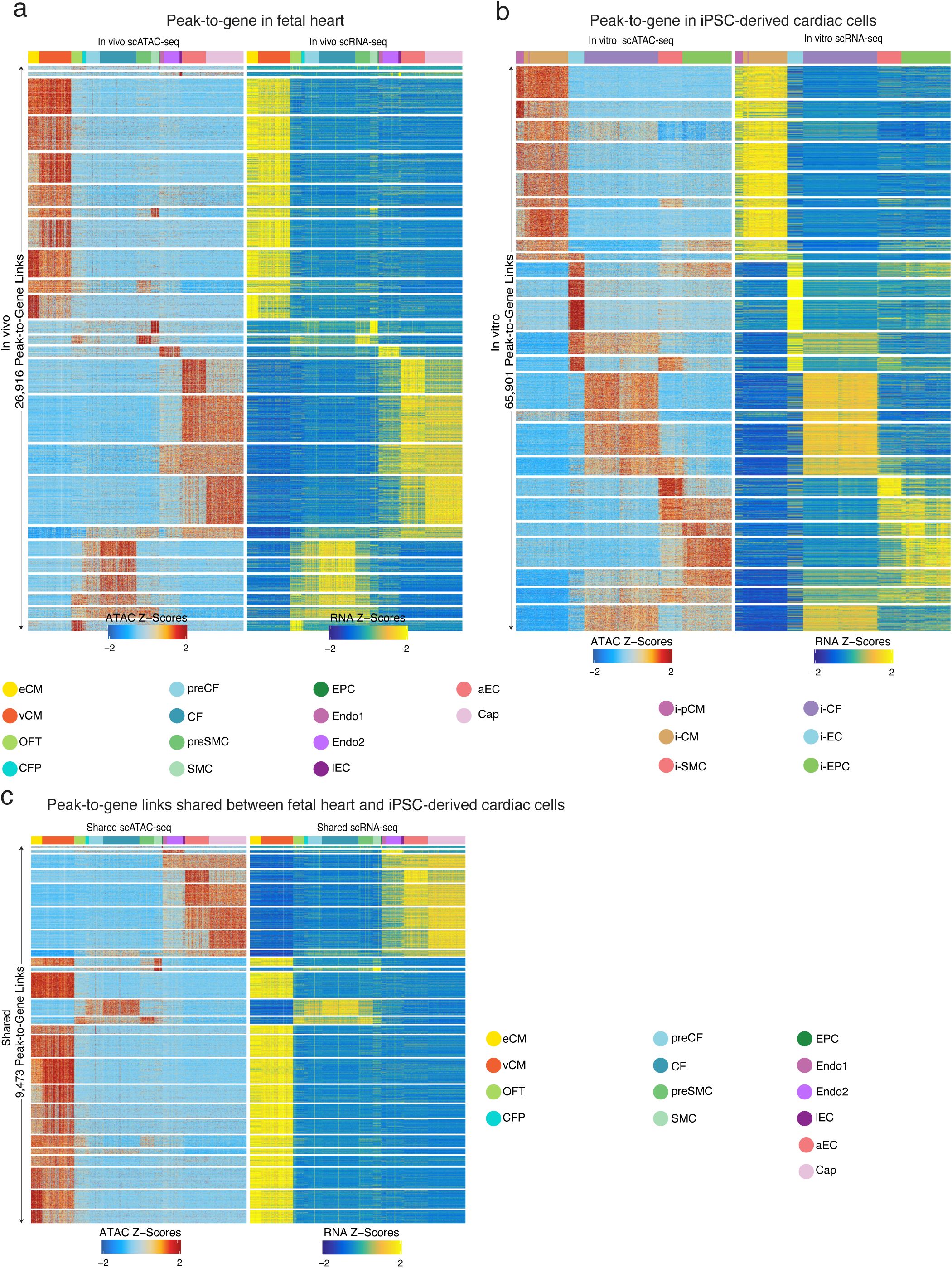
Comparison of peak to gene links between *in vivo* and *in vitro* cardiac cell types. In each panel, the left heatmap shows scATAC-seq signal (*z*-scores of log_2_(scATAC-seq read counts)) of clustered peaks across cells. The right heatmap shows the corresponding expression (*z*-scores of (log_2_(TP10K) from CCA imputed scRNA-seq) of genes linked to peaks shown in the left panel. The peaks and genes shown are based on peak-to-gene links unique to *in vivo* fetal heart cell types **(a)**, unique to fetal heart cell types (*in vivo*) **(b)**, unique to iPSC-derived cardiac cell types (*in vitro*) (**c**) and shared between *in vivo* and *in vitro* cardiac cell types.

**SFigure 15:**
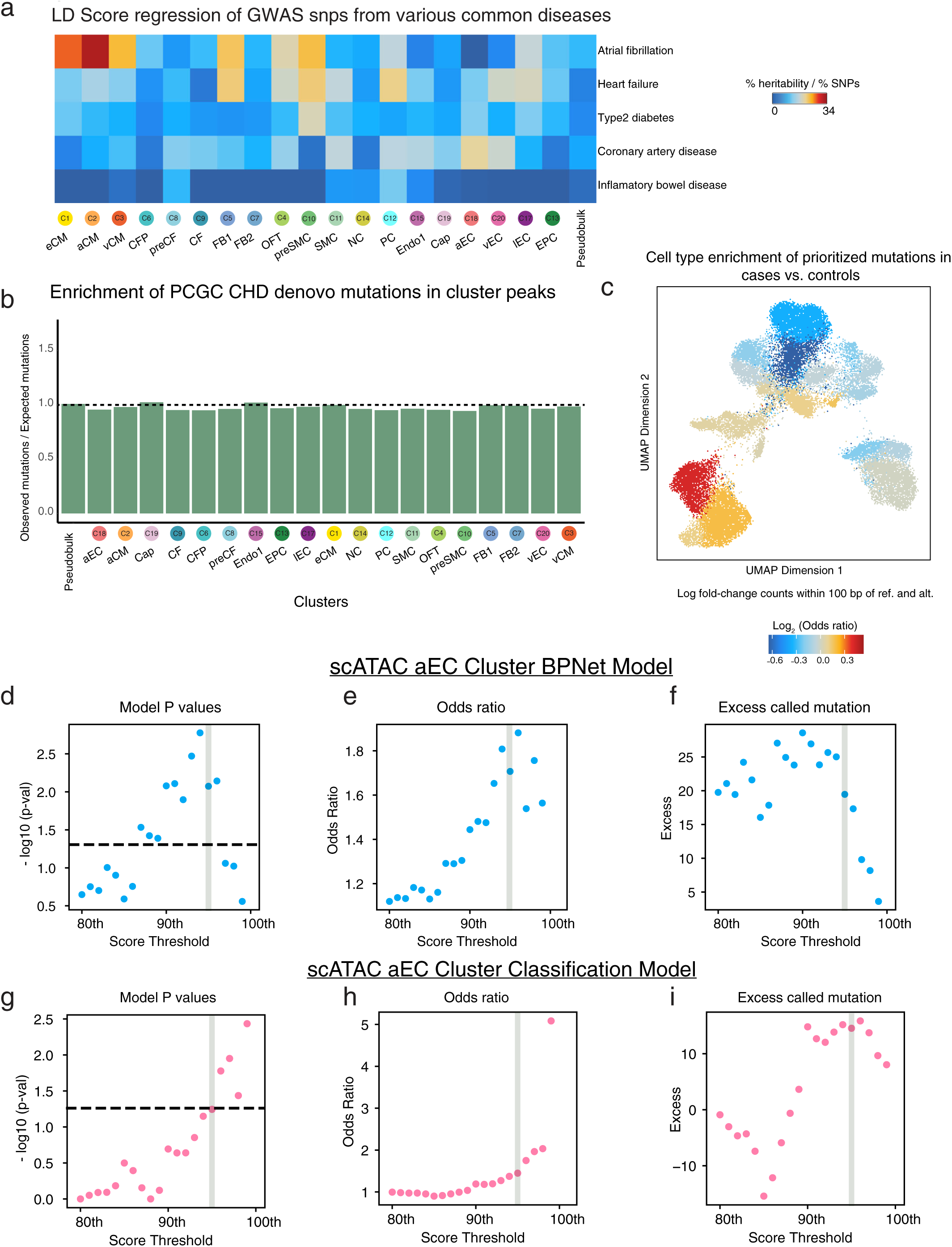
Prioritizing disease-associated non-coding variants using the cell-type resolved scATAC-seq and predictive sequence models. **(a)** Enrichment of proportion of heritability from GWAS summary statistics of cardiovascular diseases, traits and control diseases (rows) attributed to scATAC-seq derived cRE landscapes of *in vivo* and *in vitro* cardiac cell types (columns). **(b)** Enrichment of cases versus control mutations using naïve overlap with cluster-specific ATAC-seq peaks, showing relevance of the deep learning model to capture pathogenic disruptions. **(c)** Enrichment (log2(OR) counts within +/- 50 bp, Fisher’s Exact Test) of prioritized mutations from each cell-type specific BPNet model in CHD cases vs. controls plotted on the scATAC-seq UMAP of all fetal heart cells. (**d, e & f**) Evaluation of robustness in disease prioritization of aEC model across different threshold values. (**d**) the log (Fisher’s exact test p-value), (e) the Fisher’s exact test odds ratio and (**f**) excess number of causal mutations observed in cases compared to controls are plotted across all threshold values. (**g, h, i**) Similar metrics as (**d,e,f**) for a classification model with the same parameters as the BPNet model in aEC cluster..

**SFigure 16:**
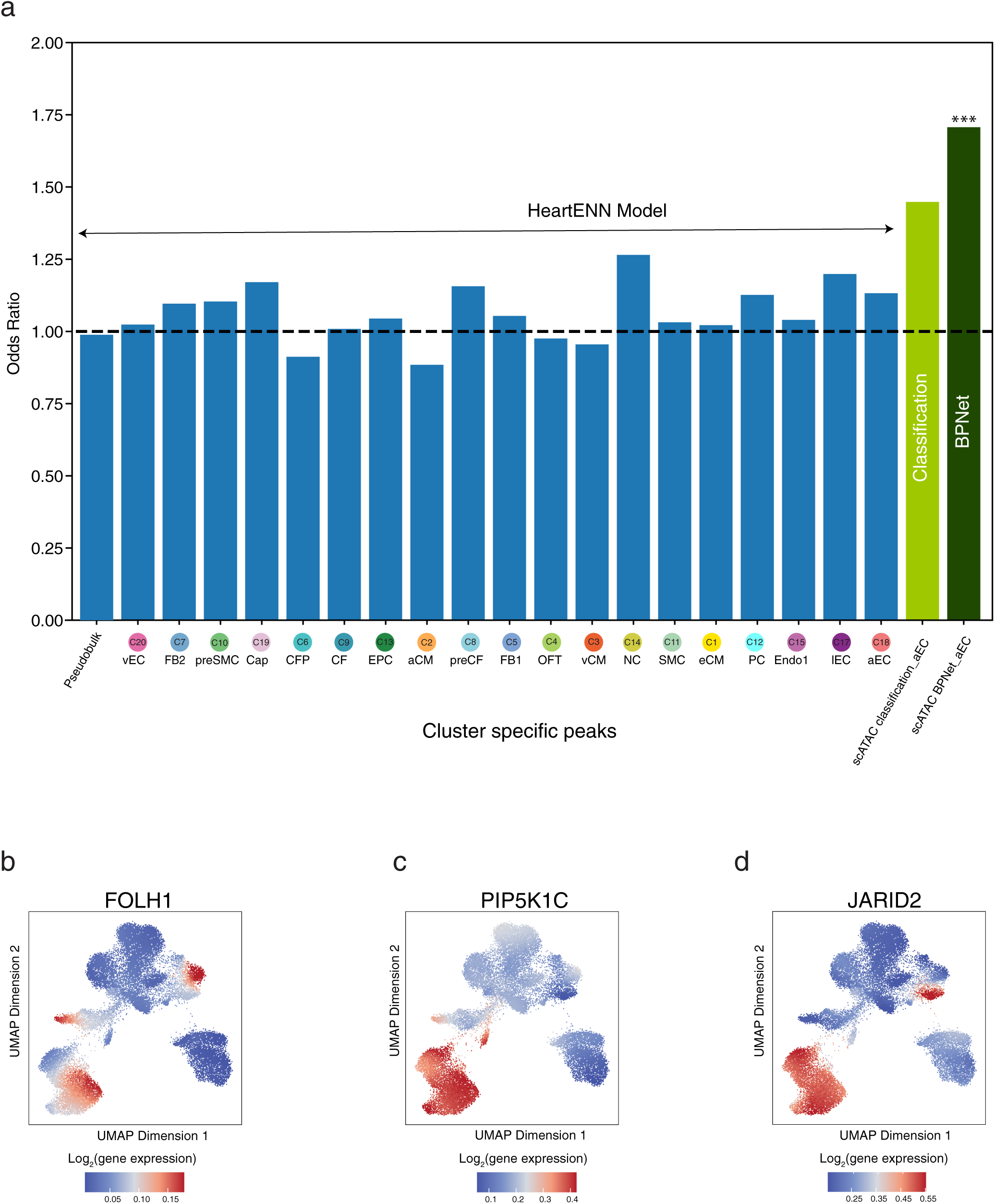
Comparison of model performance on denovo mutations: **(a)** Barplot indicating the Fisher’s exact test odds ratio of the HeartENN model (Richter et al.) subsetted to the denovo mutations in cases and controls overlapping cell type resolved peaksets (blue) scoring above 0.01 as recommended by (Richter et al) vs classification model in aEC cluster (light green) and BPNet model in aEC cluster (dark green). Stars indicate statistical significance. (* <0.05. ** =0.008). (**b, c, d**) Gene expression of *FOLH1*(**a**)*, PIP5K1C*(**b**) *& JARID2*(**c**) genes in UMAP of cells based on scATAC-seq data. Units: log_2_(TP10K).

**SFigure 17:**
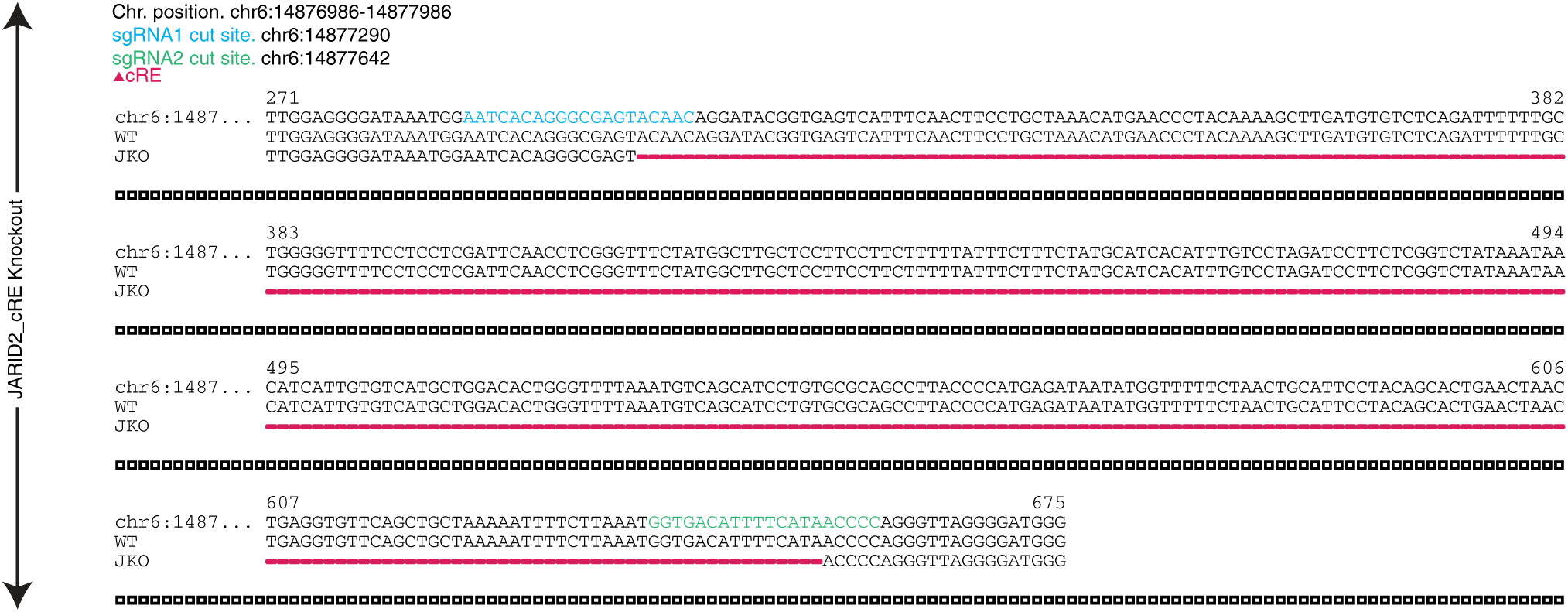
CRISPR/Cas9 deletion of JARID1 enhancer. Sanger sequencing confirms CRISPR/Cas9 targeted homozygous deletion in iPSC at the JARID2 cRE (red line).

## List of supplementary tables

TableS1 Barcode metadata for scATAC-seq experiments in fetal heart samples (post-filtering)

TableS2 Cell type annotations for all clusters identified in scATAC-seq experiments in fetal heart samples

TableS3 Marker genes (gene score) for fetal heart scATAC-seq clusters using differential test

TableS4 Gene Ontology enrichments for all scATAC-seq clusters

TableS5 Barcode metadata for scRNA-seq fetal heart samples (post filtering)

TableS6 Mapping of scRNA-seq and scATAC-seq fetal heart barcodes using CCA

TableS7 Gene expression and gene activity score correlation

TableS8 BPNet model total count prediction performance metrics

TableS9 BPNet model profile prediction performance metrics

TableS10 Pairwise co-occurence statistics of active motifs of 4 TFs highlighted in Figure 2 e,f & g panels

TableS11 TF gene expression and TF ChromVAR deviation score correlation

TableS12 Optimal transport derived attributes for each barcode

TableS13 Barcode metadata for scATAC-seq iPSC-derived cardiac cell types

TableS14 Cell type annotations for all clusters identified in scATAC-seq experiments in iPSC-derived cardiac cell types

TableS15 Marker genes (gene score) for iPSC-derived scATAC-seq clusters using differential test

TableS16 Barcode metadata for scRNA-seq from Friedman, *et al*. (post-filtering)

TableS17 Peak to gene links in fetal hearts

TableS18 Peak to gene links in iPSC-derived cardiac cell types

TableS19 Statistical significance of S-LDSC enrichments for various GWAS studies in scATAC-seq peaks from all fetal heart cell types.

TableS20 S-LDSC enrichments for various GWAS studies in scATAC-seq peaks from all fetal heart cell types.

TableS21 *De novo*, non-coding point mutations in congenital heart disease (CHD) cases from Richter *et al*.

TableS22 *De novo*, non-coding point mutations in healthy controls from Simon Simplex Collection from Richter *et al*.

TableS23 Prioritized CHD mutations using fetal heart cell-type specific BPNet models

TableS24 Prioritized control mutations using fetal heart cell-type specific BPNet models

TableS25 High confidence CHD mutations priortized in aEC

